# ComputAgeBench: Epigenetic Aging Clocks Benchmark

**DOI:** 10.1101/2024.06.06.597715

**Authors:** Dmitrii Kriukov, Evgeniy Efimov, Ekaterina Kuzmina, Anastasiia Dudkovskaia, Ekaterina E. Khrameeva, Dmitry V. Dylov

## Abstract

The success of clinical trials of longevity drugs relies heavily on identifying integrative health and aging biomarkers, such as biological age. Epigenetic aging clocks predict the biological age of an individual using their DNA methylation profiles, commonly retrieved from blood samples. However, there is no standardized methodology to validate and compare epigenetic clock models as yet. We propose ***ComputAgeBench***, a unifying framework that comprises such a methodology and a dataset for comprehensive benchmarking of different clinically relevant aging clocks. Our methodology exploits the core idea that reliable aging clocks must be able to distinguish between healthy individuals and those with aging-accelerating conditions. Specifically, we collected and harmonized 66 public datasets of blood DNA methylation, covering 19 such conditions across different ages, and tested 13 published clock models. Additionally, we compiled 46 separate datasets to facilitate the training of new aging clocks. We believe our work will bring the fields of aging biology and machine learning closer together for the research on reliable biomarkers of health and aging.

**Code:** https://github.com/ComputationalAgingLab/ComputAge

**Dataset:** https://huggingface.co/datasets/computage/computage_bench

## 1 Introduction

Longevity drugs (*a*.*k*.*a*., *geroprotectors*) appear to be on the brink of entering clinical practice to slow down or reverse the features of aging [1, 2]. The research community is yet to identify proper biomarkers of aging and rejuvenation that could be used as clinical trial endpoints instead of or in combination with observations on patient lifespans [3]. Biological age (BA) has been proposed as one of such surrogate biomarkers of aging, defined as a *generalized measure of human health* compared to the average health of individuals at a given age within a population [4, 5]. Thus, if an individual has a biological age of 40 at the chronological age of 30, it is assumed that their overall health corresponds to that of an average 40-year-old in the population. This relationship can be concisely expressed as

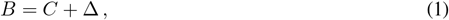

where *B* represents biological age, *C* denotes chronological age (*i*.*e*., time since birth), and Δ symbolizes BA *acceleration* (or deceleration, if negative).

In general, BA can be estimated from a set of biomarkers *X* with a model (algorithm) *f* : *X* → *B*, also called an *aging clock*. However, BA is latent: it has no ground truth value that can be measured directly and then used to train an aging clock model *f* in a classical supervised fashion, making clock validation a nontrivial task [6]. This obstacle forces researchers to introduce various additional assumptions about the aging clock behavior [7–10], as well as to experiment with different machine learning models (including penalized linear regressions, such as ElasticNet, support vector machines, decision trees, transfomer-based neural networks, *etc*. [10, 11]) and underlying types of data *X* [12– 14]. The vast majority of aging clocks, though, rely primarily on DNA methylation data, also called *epigenetic* aging clocks [15–19]. Summarizing abundant discussions about a “good” mathematical description of BA in the literature [1, 10, 20], we elicited four of its defining properties, formalized as follows.

Let *X* ∈ ℝ^*p*^, where *p* is the number of biomarkers in data, *B* ∈ ℝ, and *f* : *X* → *B*. Given the aging acceleration Δ = *B* − *C*, the following four properties hold:

1. *B* is expressed in the same time units as *C*;
2. Δ allows distinguishing between healthy individuals and individuals with aging-accelerating or decelerating conditions (AACs or ADCs), such as severe chronic diseases;
3. *B* helps to predict the remaining lifespan better than *C* does [20];
4. *B* helps to predict the time to onset of chronic age-related diseases (*e*.*g*., the Alzheimer’s) better than *C* does [20].

Garnered together, these properties motivated us to construct a benchmarking methodology for validating the potential biological age predictors. In property №1, the model *f* should output age values in a biologically meaningful range, comparable with a typical lifespan, *e*.*g*., from 0 to 120 years for the humans. This property is trivial, but necessary to differentiate biological age from other possible aging scores. To investigate if a model *f* satisfies the 2^*nd*^ property, we can define a panel of aging-accelerating (or decelerating) conditions and test if the predicted Δ allows distinguishing the individuals with an AAC/ADC from a healthy control group, according to an appropriate statistical test. To validate the compliance with the 3^*rd*^ and the 4^*th*^ properties, one also needs data on mortality and multi-morbidity. That is, the information about the timing of death or the onset of chronic age-related diseases, along with a prior measurement of a set of relevant biomarkers. It is important to note that such data are highly sensitive and are generally not publicly available.

*DNA methylation* (DNAm) is the most prevalent measurement employed in the construction of aging clocks [21]. From a chemical point of view, DNA methylation refers to a covalent modification of DNA nucleotides by the methyl groups [22]. Specifically, cytosine nucleotides (C) followed by guanine nucleotides (G), also referred to as cytosines in a CpG context or simply CpG sites (CpGs), are methylated most often in the mammalian cells, making it the most well-studied type of DNA methylation [23] (refer to Figure 1 for visualization of the DNA and CpGs). This epigenetic modification plays a crucial role in regulating gene expression and is engaged in a variety of cellular events, varying significantly across different species, tissues, and the lifespan. DNA methylation levels per site are usually reported quantitatively as beta values that represent the methylation proportion at a specific CpG site in the range from 0 to 1, where 0 indicates no methylation, and 1 indicates complete methylation across all the cells in the sample (Figure 1).

**Figure 1:**
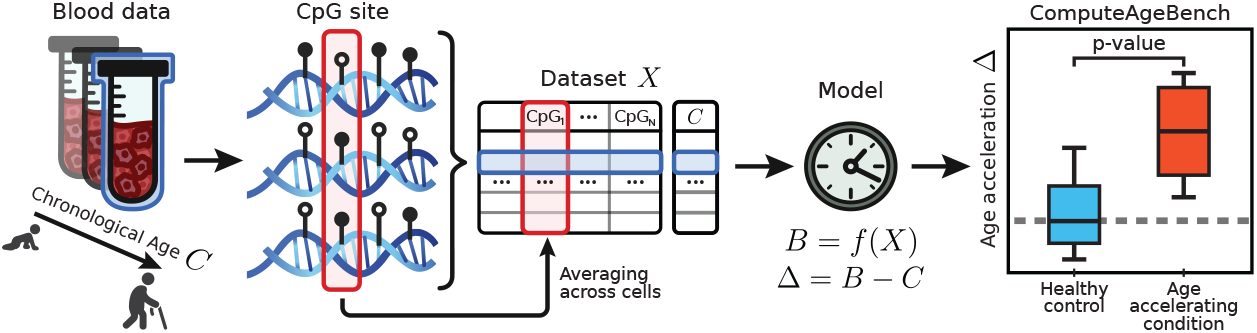
ComputAgeBench: benchmarking various epigenetic aging clock models. For a dataset *X*, obtained by profiling DNA methylation at CpG sites in bulk blood samples, an aging clock model *f* is trained to distinguish healthy individuals from those with pre-defined aging-accelerating conditions.

Importantly, despite the numerous recent publications of various aging clocks [4, 10, 21], including the ones built on DNA methylation, no systematic open-access benchmark, which would include a standardized panel of datasets, diseases, interventions, or other conditions, has been proposed to date to validate the aforementioned properties. In this paper, we introduce such a benchmark to validate the 1^*st*^ and the 2^*nd*^ properties in epigenetic aging clocks. To do this, we developed a methodology for identifying aging-accelerating conditions, which relies on simple, yet strict and evidence-based principles for defining and selecting a panel of aging-accelerating conditions. We collected an unprecedented number of DNA methylation datasets for the respective conditions from dozens of published studies. 66 of these datasets are intended for clock benchmarking, while the other 46—to facilitate the training of new epigenetic clocks. We also developed a cumulative benchmarking score that aggregates two error-based tasks and two simple, but informative tasks based on common statistical tests. Ultimately, this cumulative score enables comparing aging clock ability to satisfy the 1^*st*^ and the 2^*nd*^ properties stated above.

To demonstrate our methodology in a clinically relevant scenario, we specifically focused on the blood-, saliva-, and buccal-based epigenetic biomarkers obtained via a microarray-based technology. Such biomarkers are widespread in clinical testing and aging clock construction [10, 24]. We then examined 13 published epigenetic clocks and provided their benchmarking results.

## 2 Related work and Background

### 2.1 Aging clock construction methodology

Because the BA ground truth values cannot be measured, and, therefore, a direct validation of aging clocks is problematic, previous studies introduced various approaches to construct aging clocks with different underlying assumptions. The most widespread one, belonging to the so-called “first-generation aging clocks”, uses an assumption that a model *f* can be trained to predict chronological age, *i*.*e*., *C* = *Ĉ* + *ε* = *f* (*X*) + *ε*, and its predictions will correspond to BA: *B* = *Ĉ*. The simplicity of this approach has made it attractive for decades, and it is still used today to train new aging clocks on new types of data [25–29]. In fact, BA obtained by this approach can satisfy the 2^*nd*^ [8] and the 3^*rd*^ [30] properties from our definition. However, using this assumption in Eq. (1) leads us to the conclusion that *ε* = −Δ. It then turns out that the perfect solution of the chronological age prediction problem, *i*.*e*., minimizing the prediction error so that *ε* → 0, leads to the inability of a clock to identify any aging acceleration or deceleration. Namely, it implies that Δ → −0, which is also known as *the biomarkers paradox* [7, 31] (refer to Sluiskes et al. [6] for a more recent review). Supporting this concept, it has been shown that the clocks featuring strong correlation with the chronological age poorly correlate with the population mortality [32] (hence they fail to satisfy the 3^*rd*^ property). As a consequence, validating clock performance in terms of accuracy of chronological age prediction becomes meaningless, because high accuracy may not necessarily correspond to a biologically relevant clock. Despite the obvious methodological challenges of this approach, the vast majority of aging clocks belong to the first generation [6].

Seeking for a better solution, researchers experimented with survival models, which led to the development of “second-generation aging clocks”. In this approach, models are trained to predict time to death [16, 17, 33], and the resulting prediction is rescaled to age units to represent BA, therefore addressing the 3^*rd*^ and the 4^*th*^ properties of a “good” BA estimator. However, there is no open large-scale DNA methylation data containing time-to-death or multi-morbidity measurements, with existing studies being either available upon an authorized request or being held completely private (see Appendix A.8).

### 2.2 Attempts to compare epigenetic aging clocks

Despite reported attempts to compare the performance of different aging clocks, a benchmark with a standardized panel of datasets, diseases, interventions, or other conditions has not been proposed yet. As a result, different comparative studies employ widely varying validation data and approaches [1, 19, 30, 34–40]. As highlighted in a recent review on biomarker validation by Moqri et al. [1], “*for a reliable comparison across studies*, … *biomarker formulations should be established ‘a priori’ and not be further modified during validation”*. In the same line of thought, we propose to define a standardized and a justified procedure for clock benchmarking *before* constructing any predictive model.

Two approaches we propose as essential tasks in our benchmark entail related prior art. For example, Porter et al. [41] and Mei et al. [34] used one-sample or two-sample aging acceleration tests for clock validation. Ying et al. [19] employed two-sample tests across multiple aging clocks. These authors implicitly tested the 2^*nd*^ property of “good” aging clocks discussed above. Likewise, there were also attempts to test the 3^*rd*^ and the 4^*th*^ properties separately. In some works, including the recently updated pre-print of Biolearn [42], a Python-based framework for clock training and testing in ongoing development, authors performed Cox Proportional Hazards analysis and calculated hazard ratios with statistical significance to test if BA estimates of selected clocks are capable of predicting all-cause mortality or the onset of age-related diseases (*e*.*g*., cardiovascular events) [30, 35– 37, 40, 42]. However, these prior studies are either small-scale in terms of tested diseases [19], limited to predicting the chronological age only [38], lack standardized procedures for selecting data and conditions for clock comparison and compare only a small number of models [34, 41], or rely on mortality and disease data that are under restricted access [42]. Therefore, while developing our methodology, we attempted to mitigate all mentioned drawbacks, while balancing between framework simplicity, open source, and clinical relevance.

## 3 Benchmarking Methodology

An infographic overview of the proposed benchmarking of aging clocks is shown in Figure 2.

**Figure 2:**
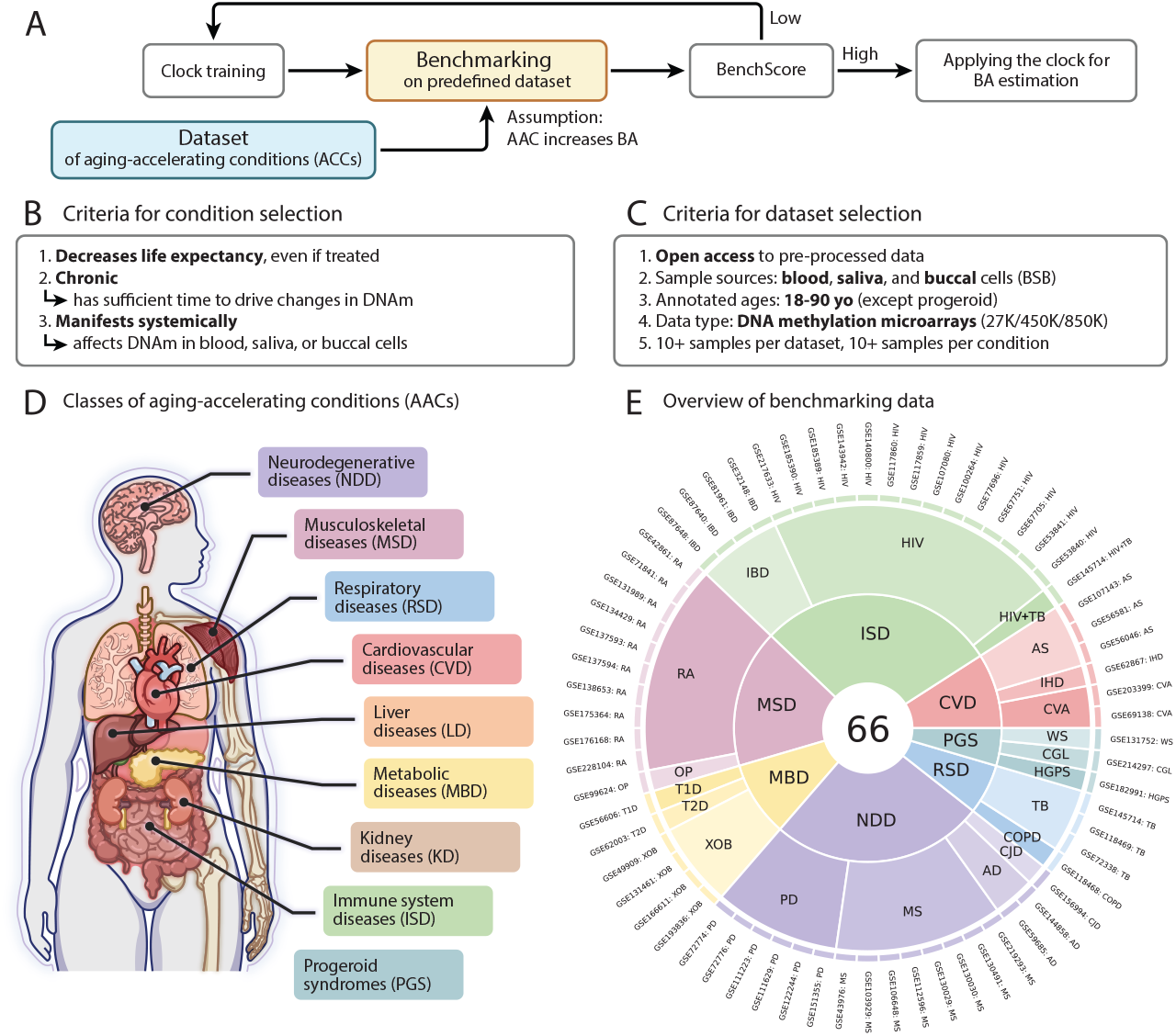
ComputAgeBench methodology. A) The proposed pipeline for constructing asging clocks features an important step of validating the model on pre-defined aging-acceleration conditions that satisfy criteria (B) and are collected into datasets that meet criteria (C) for individual study design. D) Major classes that include putative aging-accelerating conditions. E) Aggregated dataset panel for benchmarking aging clocks, comprising 66 unique data sources (labeled by their Gene Expression Omnibus dataset identification numbers and conditions) from more than 50 studies. See Table A1 for the full names and Table A2 for the population-based evidence for including each condition.

### 3.1 Criteria for selecting aging-accelerating conditions

In the context of clock benchmarking, we propose to define an aging-accelerating condition (AAC) as a biological condition that satisfies the following three criteria (Figure 2B). First, having an AAC must lead to decreased life expectancy (LE) compared to the general population, even when treated with existing therapies. Second, an AAC must be chronic (to safely assume that it has sufficient time to drive observable changes in DNAm). And third, an AAC must manifest systemically, so that it can be expected to affect DNAm in blood, saliva, and buccal cells (hereafter referred to as BSB).

Importantly, the decrease in LE and the corresponding increase in mortality must result mainly from intrinsic organismal causes rather than from socioeconomic factors and self-destructive behaviors related to a given condition. The second criterion is aimed at excluding short-term conditions such as acute infectious diseases, stressful events, and other confounding DNAm-alternating accidents, whose effects might not induce significant changes in DNAm data obtained from BSB, or, on the contrary, might last too briefly to be reliably detected. The third part of the AAC definition precludes us from considering events with long-lasting and life-threatening consequences that might be difficult to observe in BSB-derived data. For instance, a bone fracture (unless it is a critical bone marrow reserve) or some types of malignancies.

Conversely, an aging-decelerating condition (ADC) is defined as a condition that increases LE, compared to the general population, and features the same second and third criteria as an AAC. With human data, however, the ADCs are difficult to determine, as the human lifespan-increasing interventions are yet to emerge. To avoid ambiguous interpretation, we omitted such conditions in our benchmarking of human aging clocks (see Appendix A.4 and Table A1 for more details).

### 3.2 Criteria for dataset selection

Aiming to provide a comprehensive, easily accessible, and clinically relevant toolbox for the ongoing research on human epigenetic clocks, we relied on the following five criteria while performing the datasets aggregation (Figure 2C). *First*, all datasets in the benchmark must feature *open access to pre-processed data*, without any data access requests or raw data processing required. *Second*, we only used data obtained from the BSB samples. *Third*, chronological ages must be annotated with, at most, one-year intervals (*e*.*g*., without age binning by decades), including only samples from the age range of 18–90 years^1^. The only exception to this requirement are the individuals with certain *progeroid* conditions, such as the Hutchinson-Gilford progeria syndrome, who survive approximately 12 to 13 years on average: their pathologies resemble premature aging so strikingly [43] that we included patients aged under 18 years into the benchmark. *Fourth*, we employ data obtained only with the Illumina Infinium BeadChip (27K, 450K, and 850K) methylation microarrays, as they remain to be the most popular technologies for human DNAm profiling and clock construction. *Fifth*, we applied thresholds of at least 10 samples per dataset, 5 samples with an AAC per dataset, and 10 samples with an AAC across all datasets to attain sufficient statistical power.

### 3.3 Collecting AAC datasets for benchmarking

To cover as many organismal systems affected by age-related conditions as possible, we split the aggregated data into nine broad categories (Figure 2D): cardiovascular diseases (CVD), immune system diseases (ISD), kidney diseases (KD), liver diseases (LD), metabolic diseases (MBD), musculoskeletal diseases (MSD), neurodegenerative diseases (NDD), respiratory diseases (RSD), and progeroid syndromes (PGS). In each class, we identified several AACs relying on the established lists of age-related diseases and on the leading causes of death [34, 44, 45], including closely associated conditions and other conditions mentioned in a variety of epigenetic clock studies [8, 16, 19, 34, 46]. The corresponding AACs with their abbreviations and population-based evidence for their inclusion are provided in Appendix (Tables A1 and A2, respectively).

Dataset search was performed using the NCBI Gene Expression Omnibus (GEO) database, an *omics* data repository with unrestricted access (https://www.ncbi.nlm.nih.gov/geo/). We applied filters to include the *Homo sapiens* species and all types of methylation-related studies (methylation microarray data can be found in any of these study types): methylation profiling by single-nucleotide polymorphism (SNP) array, methylation profiling by array, methylation profiling by genome tiling array, and methylation profiling by high throughput sequencing.

Upon performing the dataset search, only a portion of AACs from seven condition classes were retained (see Appendix and Table A2). All five dataset selection criteria were met by none of the found kidney- and liver-related AAC datasets, and by no buccal-based dataset. The resulting list of 66 datasets [47–96] comprises 65 blood studies and 1 saliva study, and is visualized in Figure 2E. An overview of all datasets, dataset sizes, and their age distributions is provided in Figure A1. Descriptive statistics for all datasets are provided in Figure A2.

We unified the metadata of all datasets by retrieving only the relevant metadata columns and formatting them into the appropriate data types, similarly to what was proposed by Ying et al. [42]. We also added the condition and condition class annotation, thus obtaining a single metadata file with 10,410 rows (samples) and the following columns: SampleID, DatasetID (dataset GEO accession number), PlatformID (sequencing platform), Tissue (blood or saliva), CellType (whole blood or cell type after sorting), Gender, Age, Condition, and Class (see also Appendix A.10 for details on data processing).

### 3.4 Epigenetic age predictors

Any epigenetic aging clock that predicts BA in age units (or can be re-scaled to them) can be validated in our benchmark. We tested 13 publicly available epigenetic clock models trained on adult human data to evaluate sample age (Table A3), with the model coefficients retrieved from the corresponding studies. Among the collected first-generation clocks, 6 were trained purely on blood samples [15, 19, 97, 98], 3 models were trained on multiple tissues [8, 32, 46]. Among the second-generation clocks, all were blood-based, and 2 models relied entirely on CpG sites as predictive features [16, 99], while the other 2 required additional information about gender and chronological age as inputs [17, 100]. Because the clocks were trained on somewhat differing data, some clock CpGs may be lacking in the datasets outside of training, so it is an established practice to impute such missing values [42, 101, 102]. We performed this imputation by employing the “gold standard” beta values averaged for each CpG site, retrieving them from the R “SeSAMe” package [103] (for the results on comparing imputation methods, see Appendix A.7). We also ensured that no data in the benchmark was used to train any of the selected clocks, and that all clock input and output structures were consistent with each other (“harmonized”, as described by Ying et al. [42]).

### 3.5 Benchmarking tasks for evaluating aging clocks

To benchmark aging clock models, we propose four tasks: relative aging acceleration prediction (Figure 3A), absolute aging acceleration prediction (Figure 3B), chronological age prediction accuracy (Figure 3C), and systematic chronological age prediction bias (Figure 3D). In the first two tasks, the clocks are tested if they can correctly predict aging acceleration in the predefined panel of AAC datasets.

**Figure 3:**
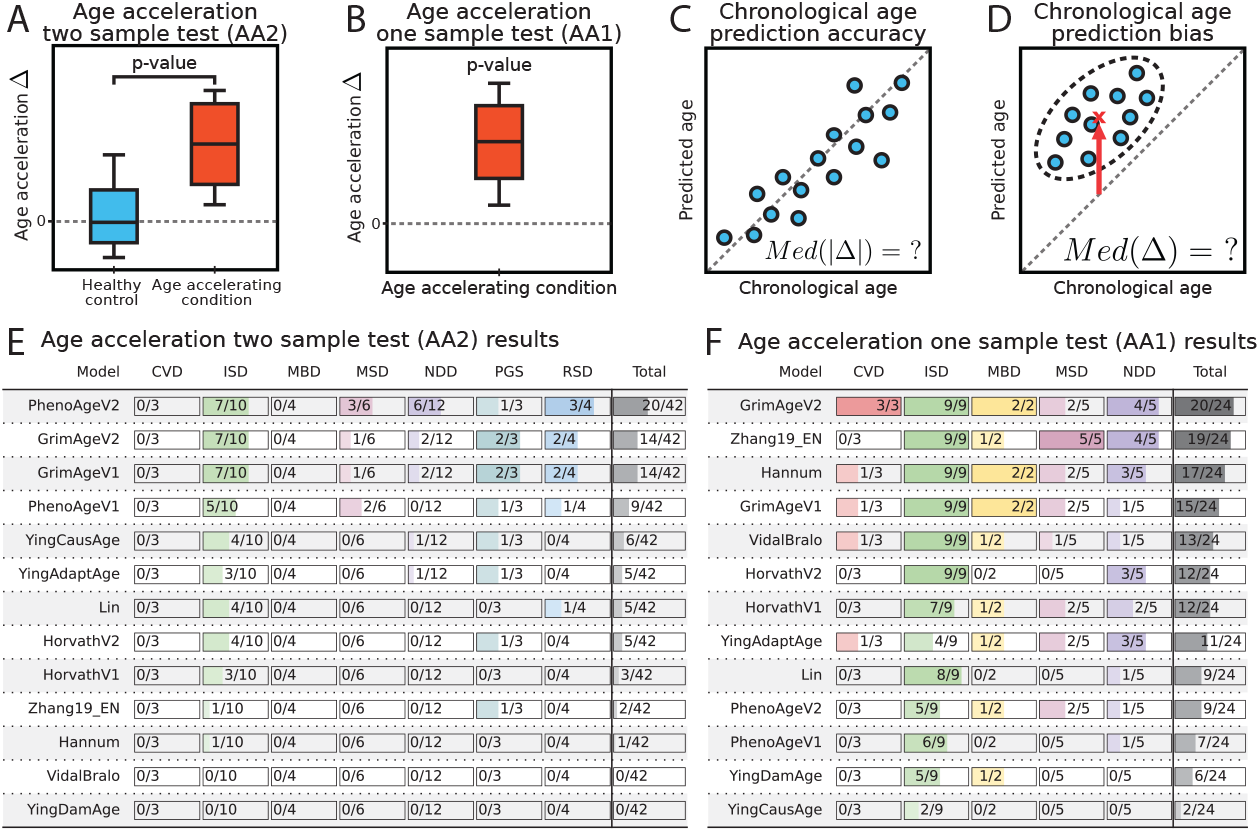
ComputAgeBench tasks and performance of aging clock models. A-D) The four bench-marking tasks. (C) illustrates that chronological age prediction accuracy is measured by median absolute error *Med*(|Δ|) across all predictions. For a limiting case of prediction bias sketched in (D), all samples were predicted with positive age acceleration, leading to a strictly positive value of *Med*(Δ), graphically represented as a red arrow pointing to a cross. E) AA2 task results split into columns by condition class, where scores demonstrate the number of datasets per class in which a given clock model detected significant difference between the HC and AAC cohorts. F) AA1 task results: same as (E), but the statistics are calculated for datasets containing the AAC cohort alone.

In the relative aging acceleration prediction task (AA2 task), we test aging clock ability to distinguish AAC from healthy control (HC) samples in a dataset containing both sample groups. After predicting ages in each dataset corresponding to this task using various clock models, we apply a two-sample Welch’s test per dataset and calculate a one-sided P-value (*i*.*e*., *H*_*A*_ : Δ_*AAC*_ *>* Δ_*HC*_) to determine if mean aging acceleration in the AAC cohort is significantly greater than that in the HC cohort (Figure 3A). Next, we apply the Benjamini-Hochberg correction procedure for controlling the false discovery rate (FDR) of predictions across all datasets, with an adjusted P-value less than 0.05 considered indicative of statistical significance. We selected a parametric test due to the assumption of normal distribution of Δ, a fundamental trait of multivariate linear regression models commonly used in aging clock construction.

In the absolute aging acceleration prediction task (AA1 task), we test clock ability to correctly predict positive aging acceleration for an AAC in the absence of the HC cohort. For each dataset in this task, we predict ages using various clock models, apply a one-sample Student’s t-test and calculate a one-sided P-value (*i*.*e*., *H*_*A*_ : Δ_*AAC*_ *>* 0) to determine if mean aging acceleration in the AAC cohort is significantly greater than zero (Figure 3B). As before, we apply the Benjamini-Hochberg correction procedure for controlling FDR with the same adjusted P-value threshold.

Clearly, the first task (AA2) provides a more rigorous way to test aging clocks compared to AA1, because it helps to control potential covariate shifts, but the second task (AA1) deserves its place in the list, as it allows introducing more data to mitigate data scarcity.

The third task is aimed at distinguishing good predictors of chronological age from predictors of biological age. Due to the paradox of biomarkers mentioned above, it is highly unlikely that the same model could combine both these properties. Yet, the good predictors of chronological age are believed to be useful in forensics [104] or data labeling, where the chronological age information is lacking. We chose median absolute aging acceleration (*Med*(|Δ|)), a full equivalent of median absolute error, for testing clock performance. We calculate it across HC samples from the whole dataset panel and report it as a single number expressed in years.

We introduced the fourth task, a prediction bias task, to evaluate the robustness of a given aging clock model to covariate shift between the original clock training dataset and the datasets from the proposed benchmark. Covariate shift, also referred to as batch effect in bioinformatics, denotes the shift between covariate distributions in two datasets. For instance, the distribution of methylation values for a given CpG site could be centered around 0.45 in one dataset and around 0.55 in the other one—a common scenario in DNAm and other omics data. Because each clock is trained on healthy controls, we expect age deviation of HC samples to be zero on average (*i*.*e*., *E*(Δ_*HC*_) = 0). In practice, however, due to the presence of a covariate shift between the training and testing data, a clock might produce biased predictions, resulting in a systemic bias and adding or subtracting extra years for a healthy individual coming from an external dataset. The goal of the fourth task is to control for such systemic bias in clock predictions. Therefore, as a benchmarking metric for this task, we calculated median aging acceleration (*Med*(Δ)) across HC samples from the entire dataset panel, which reflects the systematic shift in clock predictions caused by differences between datasets.

### 3.6 Cumulative benchmarking score

We define cumulative benchmarking score such that it would account for the main drawback of AA1 task, namely, the sensitivity to positive model bias. Let *S*_*AA*2_ denote total score of a model in AA2 task and *S*_*AA*1_ from the AA1 task (both *S*_*AA*2_ and *S*_*AA*1_ represent the number of datasets evaluated correctly by a model in the respective task), then the cumulative benchmarking score is:

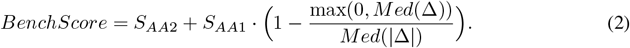

Consequently, if a model is positively biased, its performance in the AA1 task will be penalized by the bracketed coefficient by the *S*_*AA*1_, the largest when the model bias *Med*(Δ) is zero. Because *Med*(Δ) ≤ *Med*(|Δ|), this coefficient is limited to the [0, 1] interval.

While designing our metric, we aimed for simplicity and interpretability. At the same time, we sought to include more data in the benchmark to address data scarcity caused by the underrepresentation of certain AACs. Admittedly, there could be a more optimal solution for the metric, but we also believe that such a consensus solution must be proposed by a continuous collaborative discussion between the aging clock and machine learning communities, which we are eager to establish.

### 3.7 Collecting HC datasets for training

To facilitate the ML community’s engagement in developing reliable aging biomarkers, we applied our methodology to collect and unify the pre-processed datasets of HC patients, suitable for training novel aging clocks. This training data comprise 46 distinct datasets [102, 105–146] encompassing 7,419 DNA methylation profiles (see Figure A3) with annotated chronological ages. These datasets enable the training of first-generation aging clocks and the evaluation of various model architectures to determine those which best satisfy the aforementioned properties №1–4, thereby achieving superior performance on the proposed benchmark.

## 4 Results

The most rigorous of the four, AA2 task demonstrates that second-generation clocks (PhenoAgeV2 [99], GrimAgeV1 [17], GrimAgeV2 [100], and PhenoAgeV1 [16]) appear on top, particularly at predicting aging acceleration for the ISD class (Figure 3E, Supplementary Material Figure A6). Nevertheless, all clocks failed to detect aging acceleration in patients with cardiovascular and metabolic diseases, at least at the statistically significant level (see Figures A4 and A5 for results without FDR correction). Modest scores (*<*50% datasets in total) on the AA2 task across all models are expected, as no clocks had specifically been calibrated to pass this benchmarking task (and not a single first-generation clock had been trained to predict anything aside from chronological age).

In contrast, the first-generation aging clocks by Zhang et al. [32] and Hannum et al. [15] populated the top of the AA1 leaderboard, in addition to the GrimAge, exhibiting good scores across multiple condition classes (Figure 3F, Supplementary Material Figure A7). Notably, combining the results of this task with the model bias task exposes the potential source of the exceptional “robustness” in predicting accelerated aging in datasets without healthy controls.

The task of chronological age prediction accuracy reveals two undeniable leaders (Table 1): HorvathV2 [46] and HorvathV1 [8] clocks, specifically tuned for this task on large multi-tissue datasets. Notably, clocks predicting chronological age with *Med*(|Δ|) ≥ 18 years would be inferior to a constant model yielding a 50 y.o. prediction (average age across all HC samples in the benchmark). Unless scaled, such clocks can hardly be used for inferring age acceleration.

**Table 1:**
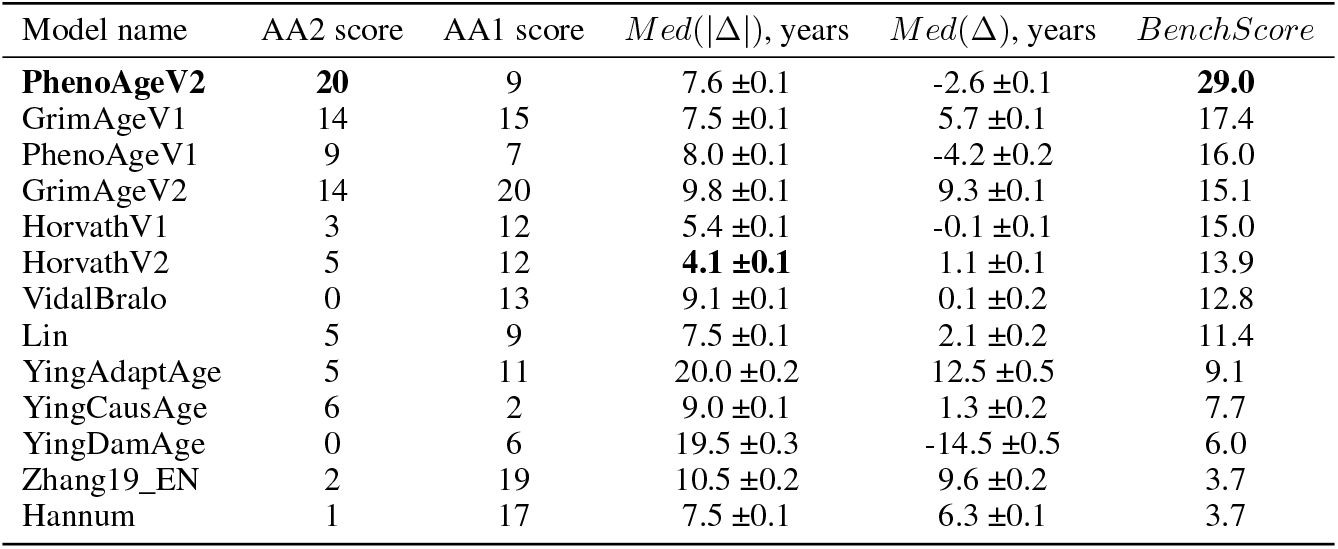
Benchmarking results.

Finally, to prove the validity of AA1 performance, a clock should also pass the task for being unbiased. We show that the AA1 leader, GrimAgeV2 clock [100], is also characterized by a large prediction bias for the HC samples (Table 1), warning us against considering its AA1 task score reliable. On the other hand, the top-2 unbiased HorvathV1 clock [8] and VidalBralo clock [98] have low prediction bias, rendering their AA1 performance as more trustworthy.

To account for the discrepancies of AA1 task interpretation regarding the prediction bias, we devised *cumulative benchmarking score* (Table 1) which penalizes AA1 score by the magnitude of prediction bias (see Eq. 2). With such a metric, a second-generation aging clock PhenoAgeV2 [99] becomes the most robust model in terms of distinguishing individuals with aging-accelerating conditions from the healthy cohort. This model is a leader, according to the cumulative benchmarking score and the AA2 task score. Closely behind it are the other second-generation clocks: GrimAgeV1 [17], PhenoAgeV1 [16], and GrimAgeV2 [100]. On the other hand, our results indicate that even the classic first-generation aging clocks, such as HorvathV1 [8] and HorvathV2 [46], can perform quite reliably in predicting biological age, at least for some condition classes. It is noteworthy that in both AA1 and AA2 tasks, many aging clocks perform well in detecting accelerated aging caused by immune system diseases, mostly represented by the human immunodeficiency virus (HIV) infection in our dataset, while the other disease classes are only captured by *some* clocks, putatively indicating that the models could have been implicitly and unintentionally trained to recognize only a certain subset of diseases. By providing a full decomposition of our benchmarking metrics in Figures 3E,F and Table 1, we strived to clarify such skewness towards particular condition classes and allow for a detailed examination of each clock’s performance. Our analysis generalizes previous findings [34] and shows that comprehensive benchmarking of aging clocks can resolve the controversy regarding their robustness and utility.

## 5 Discussion

Biological age is an elusive concept that cannot be measured and validated directly, which necessitates careful choice of model assumptions to avoid methodological errors and false discoveries while estimating it. While maintaining some degree of correlation between predicted and chronological age is desirable, the biomarkers paradox [7] precludes one from automatically considering BA evaluation via the classic performance metrics of chronological age prediction accuracy as acceptable. From a methodological perspective, training BA predictors to estimate time to death or to a disease onset remains the most rigorous approach to aging clock validation, as these events can be measured directly. However, obtaining such data is challenging due to various ethical and financial constraints. At present, no open-access data of DNA methylation annotated with either mortality or disease onset labels are available for public clock benchmarking (see Appendix A.8).

While mortality data remain unavailable, we propose to validate clocks by their ability to demonstrate BA acceleration *in a fixed pre-determined panel of datasets* for established aging-accelerating diseases or predict decelerated aging in the datasets of lifespan-prolonging interventions. For that, we developed our benchmark, where each aging clock could be tested across 4 distinct tasks. **We gathered an unprecedented number of DNA methylation datasets from 66 studies covering 19 putative aging-accelerating conditions and 46 more datasets of healthy individuals suitable for model training** to provide the ML community with a convenient entry point into the realm of aging biomarkers. Notably, no aging-decelerating conditions have been confirmed for the benchmark study (see Appendix A.4). It should be taken into account that *in vitro* cell reprogramming cannot serve as validation data for the deceleration effect, because, as has previously been shown [147], such data are essentially out-of-domain with regard to blood DNA methylation across aging.

To showcase our benchmark, we tested 13 different published models and revealed that second-generation aging clocks, namely, PhenoAge [16], GrimAge [17], and their upgraded variants [99, 100], appeared the most successful, according to the cumulative benchmarking score. As these clocks had initially been designed to predict all-cause mortality, they were expected to be robust in distinguishing aging-accelerating conditions. Yet, our findings reinforce the growing trends in training BA predictors based on mortality rather than on chronological age [1, 4].

As blood DNA methylation generally comes from the immune cells, which would be directly affected by the HIV, it is not surprising that the majority of clocks managed to discern accelerated aging in the immune system-related conditions (featured predominantly by the HIV infection in our dataset). This result supports the notion that the blood-based clocks might be implicitly attuned to such conditions, while only a few clocks are capable of successfully capturing accelerated aging in the other disease classes.

Remarkably, some datasets were evaluated incorrectly *by all models*, which may have several possible explanations apart from poor clock performance. First, a strong covariate shift between these data and the training data might impede model performance on some datasets. Second, some selected conditions might not induce accelerated aging in blood, either by itself or by the design of the original study (see Limitations in Appendix A.1). Third, the multidimensionality of aging as a biological phenomenon might not allow for correct prediction of all aging-accelerating conditions by such univariate measures as blood-based epigenetic clocks. In favor of this notion, it has recently been shown that different organ systems have different aging trajectories [148, 149], suggesting promising directions for future research outlined in Appendix A.2.

## 6 Conclusion

In this work, we developed the first systematic benchmark for evaluating epigenetic aging clocks. We believe it will help longevity researchers and data scientists to better gauge the power of existing biomarkers of aging, quantitatively assessing their role, limitations, and reliability. We anticipate that, as a result of such computational paradigm, rapid and reliable clinical trials of lifespan-extending therapies will become an attainable reality in a not-so-distant future.

## 7 Reproducibility statement

We assured the reproducibility of our pipeline by providing a Google Colab notebook (https://colab.research.google.com/drive/1_nrGMUd8oH8ADNWUPNeXHr4ZAJlZOQhm), which allows to download all datasets and benchmark all clocks considered in this article. Additionally, we provide a notebook (https://colab.research.google.com/drive/17oODr9a5nz3GVV74Grxrxl1wxQ-Y0hAr) that briefly outlines the methodology for constructing first-generation aging clocks based on our training dataset.

## Acknowledgments and Disclosure of Funding

We thank Aleksandr Vasilev for providing his guidance on industry-level Python package development, and Viktoriia Palagina for her aid in collecting evidence on life expectancy and disease datasets. We also thank Aryuna Ayusheeva and Albina Khairetdinova for their assistance with coding and retrieving clock coefficients. We would also like to express our appreciation to Leonid Peshkin for substantive discussions in the early stages of preparing this work.

## A Appendix

### A.1 Limitations

The current version of our benchmark harbors several important limitations. First, some selected conditions might not actually fulfill the suggested criteria, especially regarding their effect on blood DNA methylation, although we did our best to identify the most unambiguous ones. From the other hand, some conditions that fit our criteria might have escaped our attention. Second, the conditions are not represented uniformly, with some being featured in 10+ datasets (HIV, rheumatoid arthritis), and some present in a single dataset with few samples (ischemic heart disease, chronic obstructive pulmonary disease, congenital generalized lipodystrophy). The third limitation arises from the known issue of hidden subgroups of patients and mislabeled instances [150]. For the AAC cohorts, having hidden co-morbidities is acceptable, as they would supposedly exaggerate aging acceleration even stronger. Conversely, having severe, but unlabeled diseases in the HC cohort would likely substantially alter the findings of our benchmark. Unfortunately, we can neither expand our dataset to cover all conditions equally by mining solely the open-access data, nor explicitly confirm if all studies at hand comply with our requirements.

### A.2 Future work

We plan to further extend our benchmarking dataset by incorporating open access data of additional modalities, such as clinical biochemistry, transcriptomics, proteomics, metabolomics, etc. To overcome the aforementioned limitations, we strongly urge an open discussion on developing a panel of conditions and datasets that would serve as the gold standard for reliable and comprehensive validation of emerging biomarkers of aging. We also believe that it is important to expand the benchmark to animal models, since collecting the required data and developing preclinical biomarkers of aging in some animals is associated with fewer ethical and financial challenges. Hopefully, all these issues and developments will be addressed by the efforts of a recently established Biomarkers of Aging Consortium (https://www.agingconsortium.org/). Ultimately, the “correct” BA estimator should satisfy all four properties we defined in the Introduction. Regardless of the clock generation or data modality, reliable aging clock models must also be able to assess the uncertainty of their own predictions before being integrated in clinical trials [147, 151]. And indeed, an example of uncertainty-aware aging clocks has recently been proposed [28]. We also aim to upgrade our package to facilitate the interaction with other clock-related resources, including Biolearn [42] and pyaging [152].

### A.3 Societal impact

The obvious positive societal impact of our work is the prospect of increased active lifespan and that of healthy longevity. Our benchmarking methodology assists in determining the most accurate predictors of the biological age, which, in turn, assists in delineating the crucial biomarkers and factors that might prolong the healthy life. The potential negative impact entails the common issues emphasized when a fundamental biological problem is tackled with the AI tools. Specific to the subject of longevity are the issues of pre-mature excitement in the mass media when a certain factor is hypothesized to prolong life. A relevant fraud in the pharmaceutical industry is also plausible, if not regulated. One could also envision the depletion of resources caused by an overpopulation of the Earth, which might happen if the longevity drug is found. These negative possibilities are not expected to be sudden and could be mitigated gradually – similarly to a plethora of other benchmarking works established for solving important biological problems.

### A.4 Motivation behind including or excluding particular conditions

Our first criterion for selecting aging-accelerating conditions (AACs) was that having an AAC must lead to decreased life expectancy (LE) compared to the general population, even when treated with existing therapies. As we have mentioned earlier, this decrease in LE and the corresponding increase in mortality must result mainly from intrinsic organismal causes rather than from socioeconomic factors and self-destructive behaviors related to a given condition. Thus, while Down syndrome (DS) is associated with elevated prevalence of multiple chronic diseases [153–155], LE of DS individuals has grown dramatically by over 450% from 1960 to 2007 [156], even though no cure for DS has been developed, suggesting strong non-biological confounding factors at play. Additionally, while some authors expect DS to display accelerated epigenetic aging [8], others anticipate deceleration when applying epigenetic clocks to DS blood samples, as DS individuals are hypothesized to feature juvenile blood [34]. Schizophrenia (SCZ) is another example of a controversial condition: while we can find increased incidence of age-related comorbidities such as cardiovascular diseases, cancers, or chronic obstructive pulmonary disease [157–159], the rates of suicide and substance-induced death are also increased in people with SCZ [157]. We therefore suggest excluding such ambiguous conditions from robust clock benchmarking, as it is currently difficult to disentangle functional organismal deterioration from external and behavioral condition-related confounders and evaluate the degree to which the latter influence LE.

Regarding cancers in general, it is difficult to formulate a pre-hoc hypothesis about the directionality of epigenetic age changes. Even though we know that DNAm can be used to create signatures of various cancers, and that changes in some DNAm sites are shared between aging and cancers [160], we cannot be certain that an aging clock would indicate accelerated aging in cancerous samples, as some cancer-specific and stem cell-like features such as telomere maintenance might prompt a clock model to treat it as a marker of partial rejuvenation. In support of these considerations, epigenetic age predictions were found to exhibit no correlation with multiple TCGA cancer types [161]. To avoid possible speculation as far as possible, we recommend excluding cancer from clock benchmarking, as it is difficult to hypothesize about clock performance in such complex phenomena.

Aging-decelerating condition (ADC) is defined as a condition that increases LE compared to the general population and features the same second and third criteria as an AAC. With respect to human data, however, the ADCs are difficult to determine, as human lifespan-increasing interventions are yet to emerge. There are genetic mutations, such as Laron syndrome (growth hormone insensitivity) or isolated growth hormone deficiency (growth hormone releasing hormone insensitivity), that appear to protect against some age-related pathologies, but they do not feature a prolonged lifespan [162]. To avoid dubious interpretation, we recommend omitting the inclusion of any condition into the ADC category when benchmarking human aging clocks.

The resulting list of condition classes and conditions selected to represent accelerated aging is listed in Table A1. Population-based evidence for condition inclusion and the number of datasets found and selected per condition are displayed in Table A2.

### A.5 On data types used for aging clocks construction

Multiple data modalities were previously used for aging clocks construction. Some examples beyond DNA methylation data include also clinical blood samples [12], psycho-social questionnaires [264], facial images [13], urine metabolites [33], and different omics data, gene expression [14], DNA accessibility [265], plasma proteins [266], etc. Interestingly, DNA methylation data allow one the most accurate prediction of chronological age compared to other data modalities, second only to facial imaging data [21], and it continues to be used most widely in aging clock construction [10]. It is also important to note that from a practical point of view, in order to construct a clinically relevant aging clock, the method of obtaining the data should not be too invasive and heavy-handed. For this reason, many clock developers prefer using blood, saliva, or buccal epithelial samples as data sources.

### A.6 Aging clocks included in the benchmarking

The full list of published aging clocks used in this analysis is provided in Table A3.

### A.7 Comparison of different approaches to missing values imputation

We ran additional experiments (see Table A4) to test different imputation methods and observed that the method we used (SeSAMe 450k) leads to the most accurate age predictions across all models except the VidalBralo clock, whose MAE is 0.19% lower when using imputation by zeros. Notably, we did not have to impute all 800k+ sites in the whole dataset, as we only imputed sites included in each respective clock model. Most importantly, clock performance in other benchmarking tasks remained intact, regardless of the imputation strategies.

**Table A1:** Aging-accelerating conditions. ICD-10: class or condition code(s) from the International Classification of Diseases Version 10; a dash indicates lack of specific code; abbr.: abbreviation.

| Condition class | Class ICD-10 code | Class abbr. | Aging-accelerating condition (AAC) | Condition ICD-10 code | Condition abbr. |
| --- | --- | --- | --- | --- | --- |
| Cardio-vascular diseases | I00-I99 | CVD | Atherosclerosis | I70 | AS |
|  |  |  | Ischemic heart disease | I20-I25 | IHD |
|  |  |  | Cerebrovascular accident | I60-I63 | CVA |
|  |  |  | Heart failure | I50 | HF |
|  |  |  | Myocardial infarction | I21-I22 | MCI |
| Immune system diseases | — | ISD | Inflammatory bowel disease | K50-K51 | IBD |
|  |  |  | Human immunodeficiency virus infection | B20-B24 | HIV |
| Kidney diseases | N00-N99 | KDD | Chronic kidney disease | N18 | CKD |
| Liver diseases | K70-K77 | LVD | Nonalcoholic steatohepatitis | K75.81 | NASH |
|  |  |  | Primary biliary cholangitis | K74.3 | PBC |
|  |  |  | Primary sclerosing cholangitis | K83.01 | PSC |
|  |  |  | Cirrhosis | K70.3, K74.3-K74.6 | CIR |
| Metabolic diseases | E00-E90 | MBD | Extreme obesity | E66.01, E66.2 | XOB |
|  |  |  | Type 1 diabetes | E10 | T1D |
|  |  |  | Type 2 diabetes | E11 | T2D |
|  |  |  | Metabolic syndrome | E88.810 | MBS |
| Musculo-skeletal diseases | M00-M99 | MSD | Sarcopenia | M62.84 | SP |
|  |  |  | Osteoporosis | M80-M81 | OP |
|  |  |  | Osteoarthritis | M15-M19 | OA |
|  |  |  | Rheumatoid arthritis | M05-M06 | RA |
| Neuro-degenerative diseases | G00-G99 | NDD | Alzheimer’s disease | G30 | AD |
|  |  |  | Parkinson’s disease | G20 | PD |
|  |  |  | Multiple sclerosis | G35 | MS |
|  |  |  | Dementia with Lewy bodies | G31.83 | DLB |
|  |  |  | Creutzfeldt-Jakob disease | A81.0 | CJD |
| Respiratory diseases | J00-J99 | RSD | Chronic obstructive pulmonary disease | J44 | COPD |
|  |  |  | Idiopathic pulmonary fibrosis | J84.112 | IPF |
|  |  |  | Tuberculosis | A15 | TB |
| Progeroid syndromes | — | PGS | Werner syndrome | E34.8 | WS |
|  |  |  | Hutchinson-Gilford progeria syndrome | E34.8 | HGPS |
|  |  |  | Congenital generalized lipodystrophy | E88.1 | CGL |
|  |  |  | Dyskeratosis congenita | Q82.8 | DKC |

**Table A2:** Population-based evidence for condition inclusion, and the number of datasets found and selected for each condition. GEO: Gene Expression Omnibus database; abbr.: abbreviation.

| **Class**<br>**abbr.** | **Condition**<br>**abbr.** | **Evidence of decreased life expectancy**<br>**and/or systemic effects** | **N items**<br>**in the**<br>**GEO query** | **N datasets**<br>**after**<br>**filtering** |
| --- | --- | --- | --- | --- |
| CVD | AS | [163–167] | 22 | 3 |
|  | IHD | [168–171] | 21 | 1 |
|  | CVA | [168, 172–174] | 10 | 2 |
|  | HF | [168, 175, 176] | 14 | 0 |
|  | MCI | [168, 177, 178] | 19 | 0 |
| ISD | IBD | [179–185] | 30 | 4 |
|  | HIV | [168, 186–189] | 44 | 15 |
| KDD | CKD | [190–192] | 6 | 0 |
| LVD | NASH | [193, 194] | 8 | 0 |
|  | PBC | [195, 196] | 1 | 0 |
|  | PSC | [197, 198] | 2 | 0 |
|  | CIR | [168, 199, 200] | 68 | 0 |
| MBD | XOB | [168, 201–204] | 96 | 4 |
|  | T1D | [205–210] | 14 | 1 |
|  | T2D | [168, 211–214] | 45 | 1 |
|  | MBS | [168, 215–217] | 17 | 0 |
| MSD | SP | [218–220] | 2 | 0 |
|  | OP | [221–224] | 5 | 1 |
|  | OA | [168, 225–227] | 26 | 0 |
|  | RA | [228–231] | 37 | 10 |
| NDD | AD | [168, 232–235] | 43 | 2 |
|  | PD | [236–238] | 37 | 6 |
|  | MS | [239–241] | 29 | 8 |
|  | DLB | [242–244] | 5 | 0 |
|  | CJD | [245–247] | 1 | 1 |
| RSD | COPD | [168, 248–250] | 14 | 1 |
|  | IPF | [251–254] | 14 | 0 |
|  | TB | [168, 255, 256] | 13 | 3 |
| PGS | WS | [257–259] | 7 | 1 |
|  | HGPS | [257, 260] | 14 | 1 |
|  | CGL | [261, 262] | 1 | 1 |
|  | DKC | [263] | 2 | 0 |
| **Total number of datasets** |  |  | **667** | **66** |

**Table A3:** Aging clock models tested in our benchmark.

| Model name | N CpGs | Generation | Extra parameters | Tissues used for training | Reference |
| --- | --- | --- | --- | --- | --- |
| Hannum | 71 | 1 | — | Blood | [15] |
| HorvathV1 | 353 | 1 | — | Multi-tissue | [8] |
| Lin | 99 | 1 | — | Blood | [97] |
| VidalBralo | 8 | 1 | — | Blood | [98] |
| HorvathV2 | 391 | 1 | — | Blood, Skin | [46] |
| PhenoAgeV1 | 513 | 2 | — | Blood | [16] |
| Zhang19_EN | 514 | 1 | — | Blood, Saliva | [32] |
| GrimAgeV1 | 1030 | 2 | Age, Sex | Blood | [17] |
| GrimAgeV2 | 1030 | 2 | Age, Sex | Blood | [100] |
| PhenoAgeV2 | 959 | 2 | — | Blood | [99] |
| YingAdaptAge | 999 | 1 | — | Blood | [19] |
| YingCausAge | 585 | 1 | — | Blood | [19] |
| YingDamAge | 1089 | 1 | — | Blood | [19] |

**Table A4:** MAE results (in years) for different strategies of missing values imputation (sorted by increasing MAE when using SeSAMe 450k imputation).

| Model name | SeSAMe 450k | Average | Zeros |
| --- | --- | --- | --- |
| HorvathV2 | **4.143847** | 4.701762 | 4.719477 |
| HorvathV1 | **5.350622** | 5.475857 | 5.475857 |
| GrimAgeV1 | **7.462245** | 8.102066 | 8.241653 |
| Lin | **7.467559** | 8.367630 | 8.429655 |
| Hannum | **7.477633** | 7.890421 | 7.907489 |
| PhenoAgeV2 | **7.604413** | 8.439977 | 8.432397 |
| PhenoAgeV1 | **8.009677** | 8.380239 | 8.381561 |
| YingCausAge | **8.969959** | 11.599078 | 11.551690 |
| VidalBralo | 9.124225 | 9.124387 | **9.107015** |
| GrimAgeV2 | **9.796544** | 10.513198 | 10.576180 |
| Zhang19_EN | **10.534452** | 10.611938 | 10.611938 |
| YingDamAge | **19.534224** | 20.179561 | 20.211066 |
| YingAdaptAge | **19.972273** | 23.287544 | 23.353844 |

### A.8 On accessibility of existing epigenetic mortality data

Although there are some existing biobanks that aggregate sensitive human data and provide them in an open-access manner, (e.g., NHANES: https://wwwn.cdc.gov/nchs/nhanes/), most biobanks rely on authorized access to their data (e.g., UK Biobank: https://www.ukbiobank.ac.uk/). The similar semi-open situation occurs with DNA methylation data. Here, we provide information about 12 cohort studies containing DNA methylation data and mortality/morbidity information simultaneously, but all of which allow downloading their data upon a reasonable request by contacting with the principal investigators of each cohort or by requesting data on a special platform. These studies include the Framingham Heart Study (FHS), the Women’s Health Initiative (WHI), the Lothian Birth Cohorts (LBC), the Atherosclerosis Risk in Communities (ARIC), the Cardiovascular Health Study (CHS), the Normative Aging Study (NAS), the Invecchiare in Chianti (InCHIANTi), the Cooperative Health Research in the Region of Augsburg (KORA), the Epidemiologische Studie zu Chancen der Verhütung, Früherkennung und optimierten Therapie chronischer Erkrankungen in der älteren Bevölkerung (ESTHER), the Danish Twin Register sample (DTR), the Rotterdam Study (RS), and the Coronary Artery Risk Development in Young Adults (CARDIA) [1, 36]. While we recognize the risks associated with releasing sensitive patient data into the public domain, we also want to emphasize that comprehensive independent validation of the aging clock is difficult without these important datasets. The confidentiality of such data also does not allow us to use it as part of this open-access benchmark. Instead, we focused solely on those epigenetic datasets of patients with AACs distributed across human lifespan, which did not contain information on mortality, but was publicly accessible.

### A.9 DNA methylation data collection

As we have mentioned in the Methodology section, dataset search was performed using the NCBI Gene Expression Omnibus (GEO) database, an unrestricted-access omics data repository (https://www.ncbi.nlm.nih.gov/geo/) which shares data using the Open Database License (ODbL). The resulting list of 66 AAC datasets [47–96] indicated in Table A2 is visualized in Figure 2E and includes: atherosclerosis (AS), ischemic heart disease (IHD, also known as coronary heart disease), cerebrovascular accident (CVA, also known as stroke), inflammatory bowel disease (IBD, including Crohn’s disease and ulcerative colitis), human immunodeficiency virus infection (HIV), extreme obesity (XOB, defined by having BMI ≥ 40 kg/m^2^ [267, 268]; also known as class III obesity, severe obesity, or morbid obesity), type 1 diabetes mellitus (T1D), type 2 diabetes mellitus (T2D), rheumatoid arthritis (RA), osteoporosis (OP), Alzheimer’s disease (AD), Parkinson’s disease (PD), multiple sclerosis (MS), Creutzfeldt-Jakob disease (CJD), chronic obstructive pulmonary disease (COPD), tuberculosis (TB), Werner syndrome (WS, including atypical Werner syndrome), Hutchinson-Gilford progeria syndrome (HGPS, including non-classical progeroid laminopathies), and congenital generalized lipodystrophy (CGL, also known as Berardinelli-Seip lipodystrophy). Age distribution across conditions is demonstrated in Figure A1. Overviews of all benchmarking and training (healthy controls only) datasets and their age distributions are provided in Figure A2 and Figure A3. The information on how patient consent was obtained and which ethics procedures were implemented can be accessed in the respective publications. As per NCBI GEO guidelines, all submitters must “ensure that the submitted information does not compromise participant privacy” (https://www.ncbi.nlm.nih.gov/geo/info/faq.html).

### A.10 DNA methylation data processing

After pre-processing raw output from microarrays or sequencing machines, DNA methylation levels per site are reported quantitatively either as beta values, or as M values. Briefly, beta values represent the ratio of methylated signal (probe intensity or sequencing read counts) to total signal per site (sum of methylated and unmethylated probe intensities or sequencing read counts), while M value is the log2 ratio of the methylated signal versus an unmethylated signal. A more thorough comparison of the two measures can be found in Du et al. [269]. In the original datasets deposited on GEO, DNA methylation values were represented either as a beta fraction (ranging from 0 to 1), beta percentages (ranging from 0 to 100), or M-values (can be both negative and positive, equals 0 when beta equals 0.5). We converted all data to the beta-value fractions ranging from 0 to 1. The values outside this range were treated as missing values (NaNs), as they are not biological. In each dataset, only samples that were relevant for benchmarking (that is, were annotated by age, tissue, and condition) were retained.

**Figure A1:**
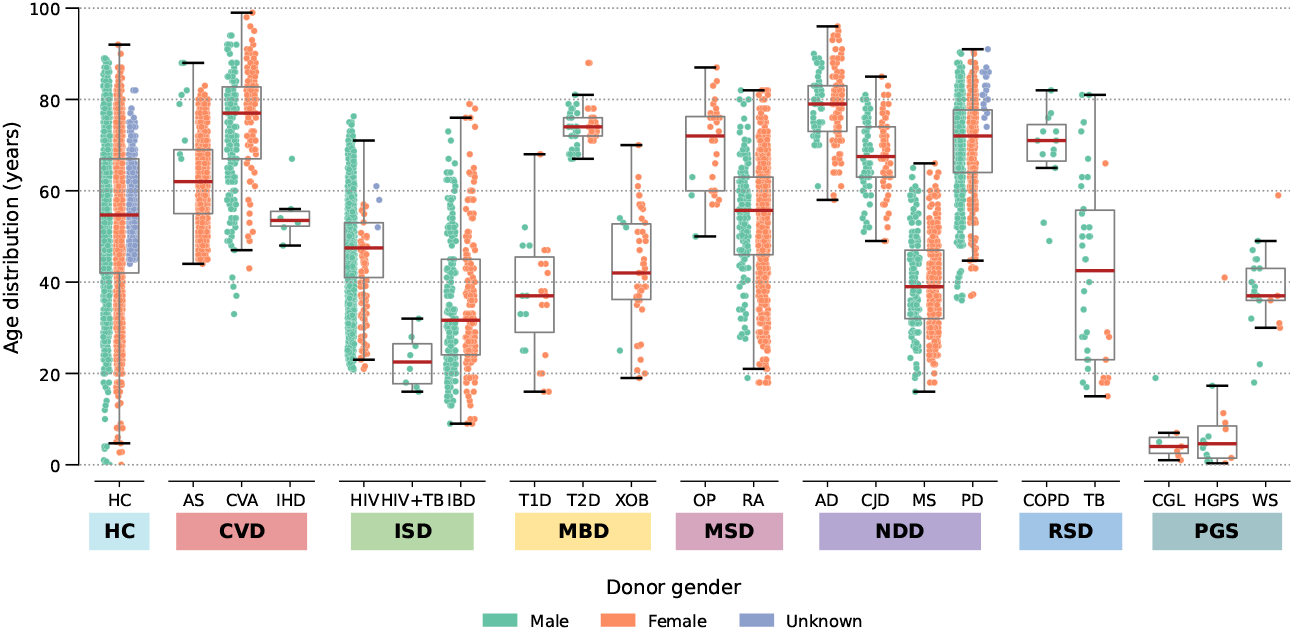
Distribution of the benchmarking dataset samples per condition across ages.

The resulting datasets meta-data contains the following fields: DatasetID (datasets GEO ID), PlatformID (GEO ID of a DNA methylation profiling platform), Tissue (sample source tissue: “Blood” stands for peripheral blood samples, “Saliva”—for saliva samples, and “Buccal”—for buccal swab samples), CellType (sample cell type: either a specific cell population, e.g., immune cell subtypes with cell type-specific molecular markers, or broader categories such as whole blood, buffy coat, peripheral blood mononuclear cells (PBMC), or peripheral blood leukocytes (PBL); some samples lack this annotation), Gender (abbreviated sample donor gender: M = Male, F = Female, U = Unknown), Age (sample donor chronological age in years; in the original datasets deposited on GEO, it can be either rounded by the researchers to full years, or converted from months, weeks, or days; where available, we calculated years from the smaller units), Condition (one of AACs or HC), and Class.

As there is no gold standard for DNAm processing, each research group carries out their preferred pipeline that does not necessarily match the processing pipeline used for training the clock model, especially in case of applying earlier clocks (e.g., those by Hannum et al. [15] or Horvath [8]) to recently collected data. Therefore, so as to retain this typical workflow and not to put any clock model into advantage by choosing the same processing that matches its own pipeline for every dataset, we did not perform any post-processing, inter-dataset normalization, or batch effect correction. In doing so, we also relied on two existing papers. First, compiling already pre-processed datasets without performing the same processing for all of them was done by Ying et al. [42] in creating Biolearn, another notable effort in the aging clock community. Second, we were encouraged by a recent work by Varshavsky et al. [28] who managed to create an accurate clock model by combining several blood datasets—without any additional normalization or correction procedure, using already pre-processed data from previous studies (some of which are included in our dataset as well), and thus demonstrating that the between-dataset normalization is not critical for this type of data.

### A.11 Benchmarking results without FDR correction

Figures A4 and A5 demonstrate benchmarking results before applying FDR correction.

**Figure A2:**
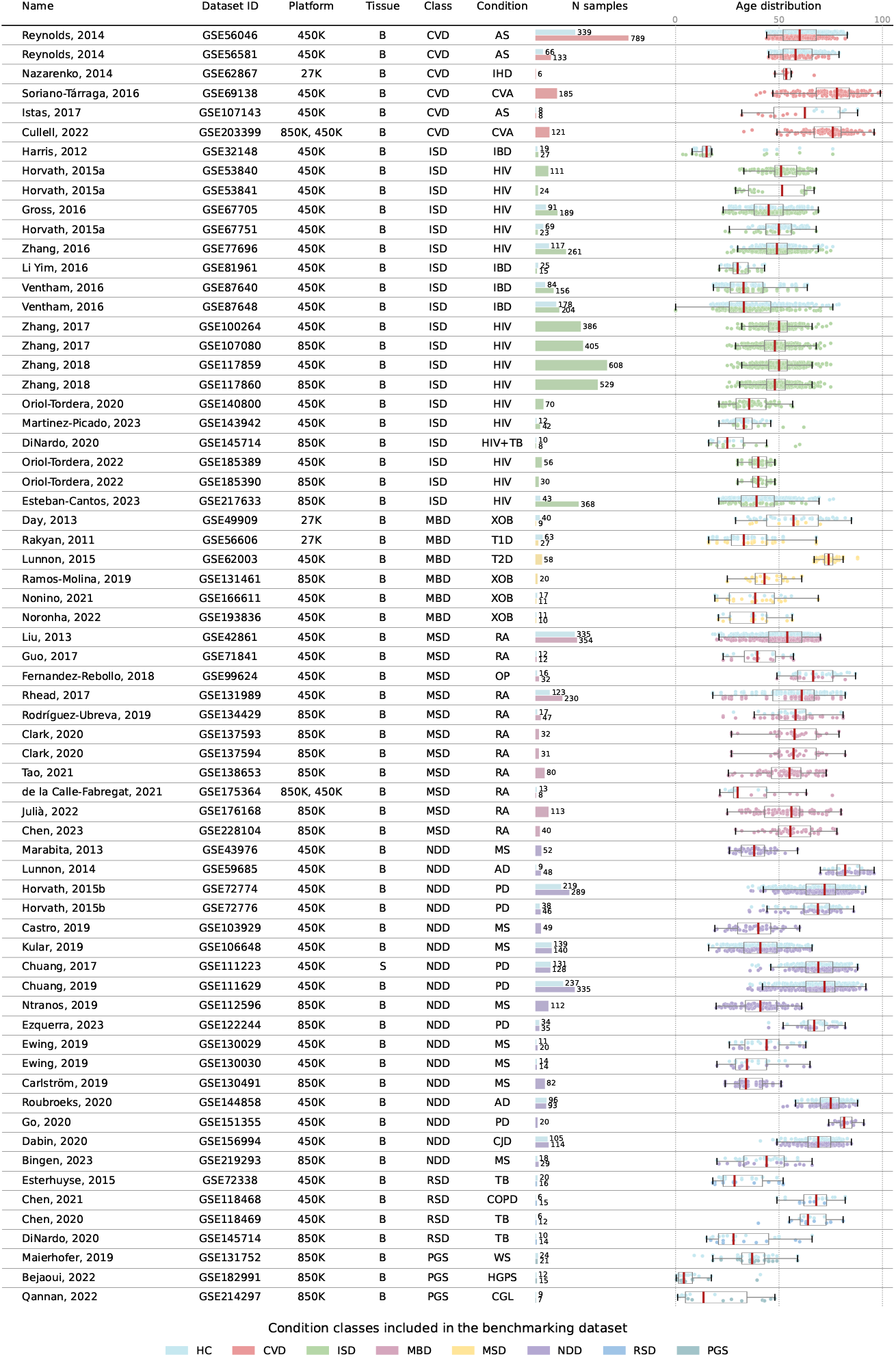
Descriptive statistics of datasets included in the benchmark. B: blood, S: saliva. Ages are indicated in years.

**Figure A3:**
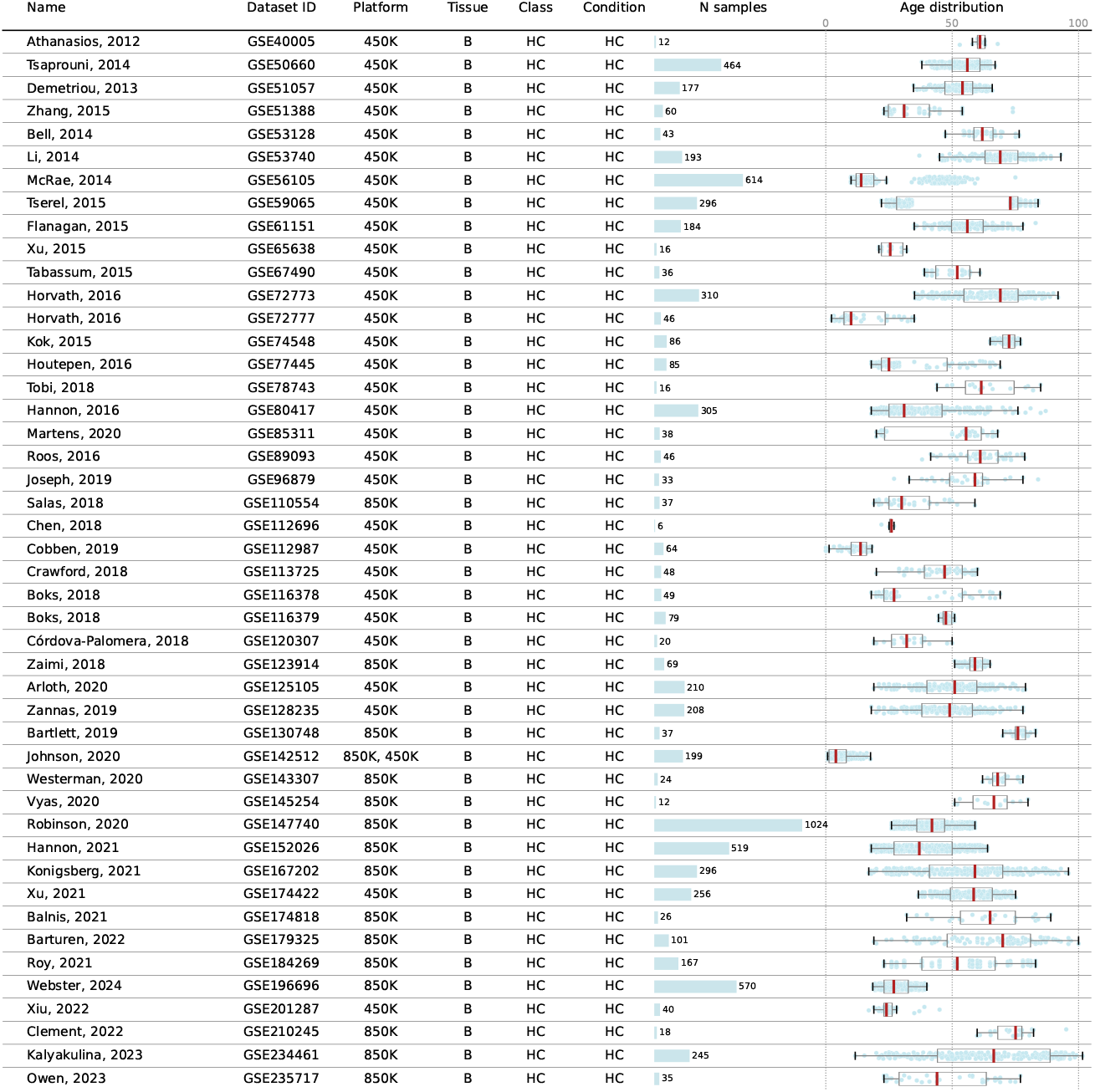
Descriptive statistics of datasets included in the training subset. B: blood. Ages are indicated in years.

**Figure A4:**
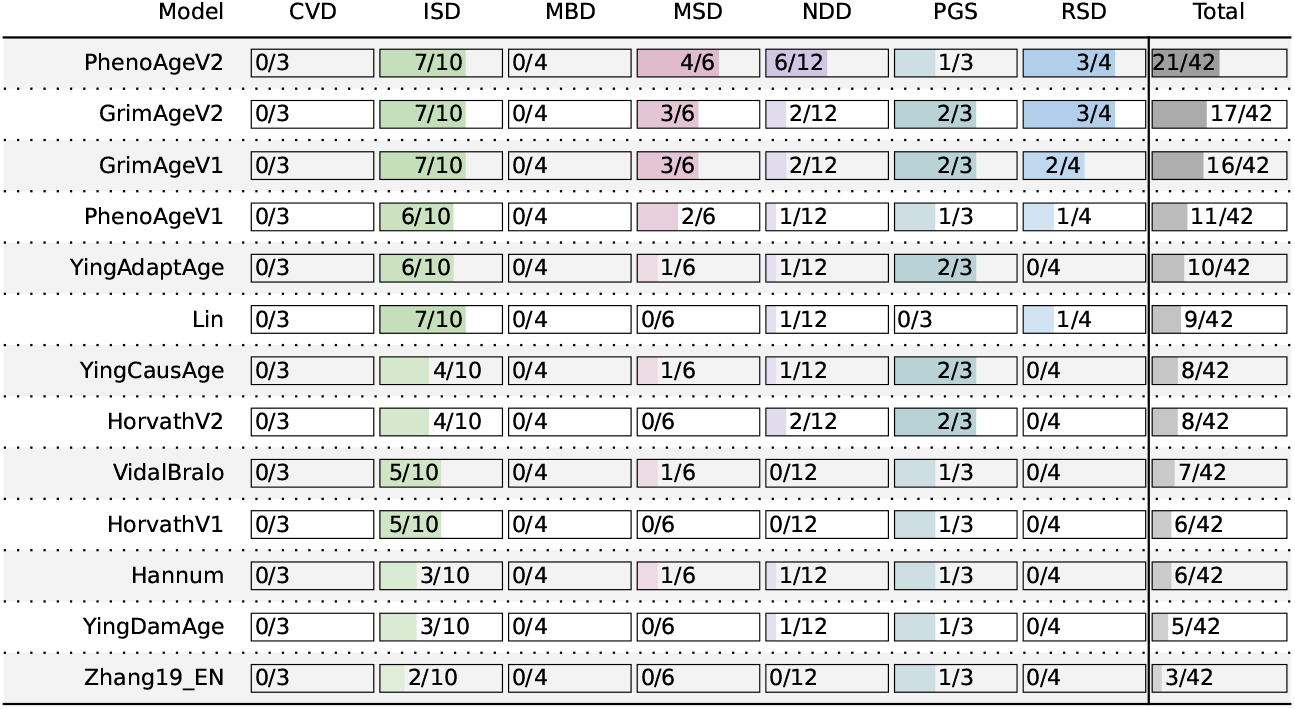
AA2 task results split into columns by condition class **without FDR correction of P-values**. Scores demonstrate the number of datasets per class, in which a given clock model detected significant (at the 0.05 level of significance) difference between the HC and AAC cohorts.

**Figure A5:**
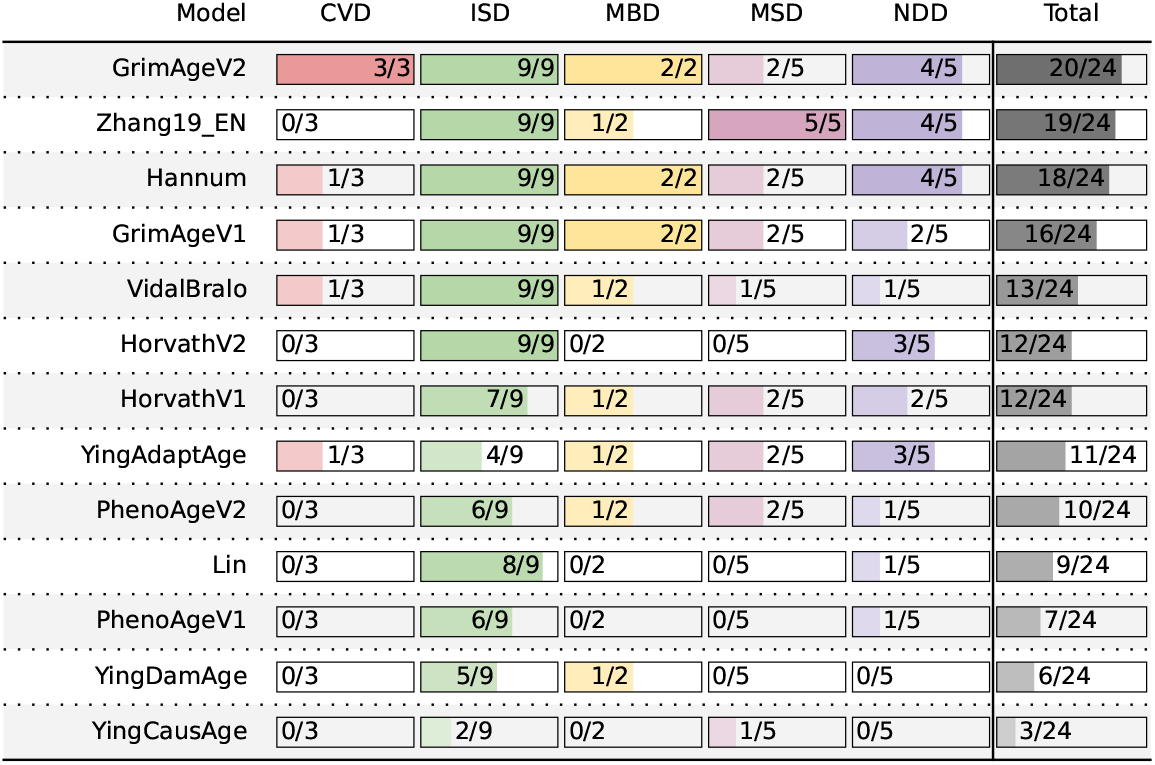
AA1 task results **without FDR correction of P-values**: same as Figure A4, but the statistics are calculated for datasets containing the AAC cohort only.

## Footnotes

1 Reporting increased or decreased biological age for people outside of this range is debatable.

