## Supplementary figures and images for "ComputAgeBench: Epigenetic Aging Clocks Benchmark"

### Figure_A6_AA2-CVD.pdf

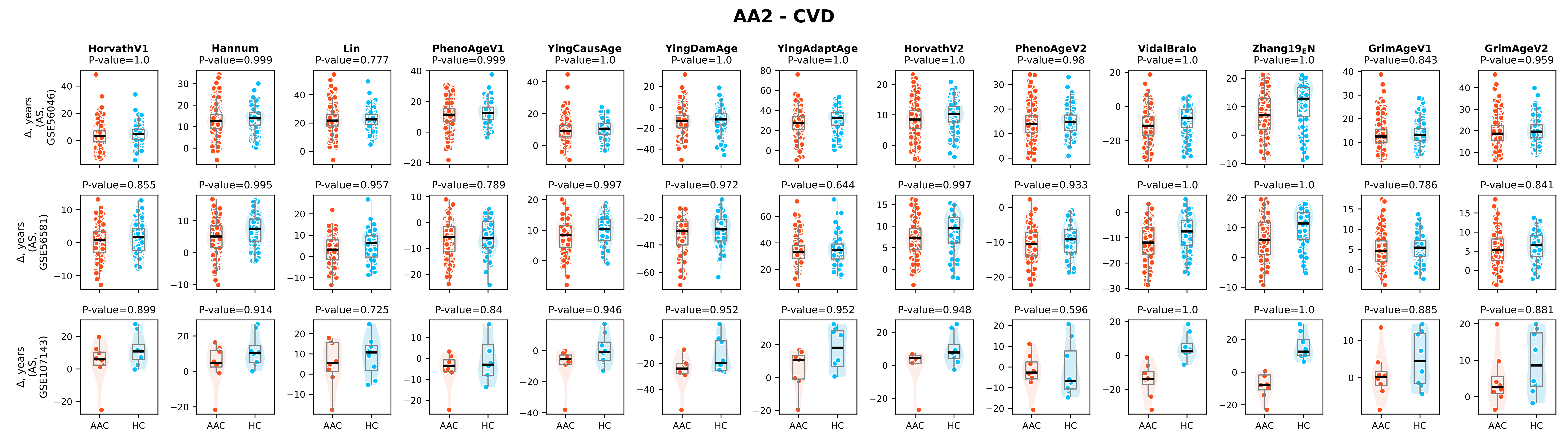

### Figure_A6_AA2-ISD.pdf

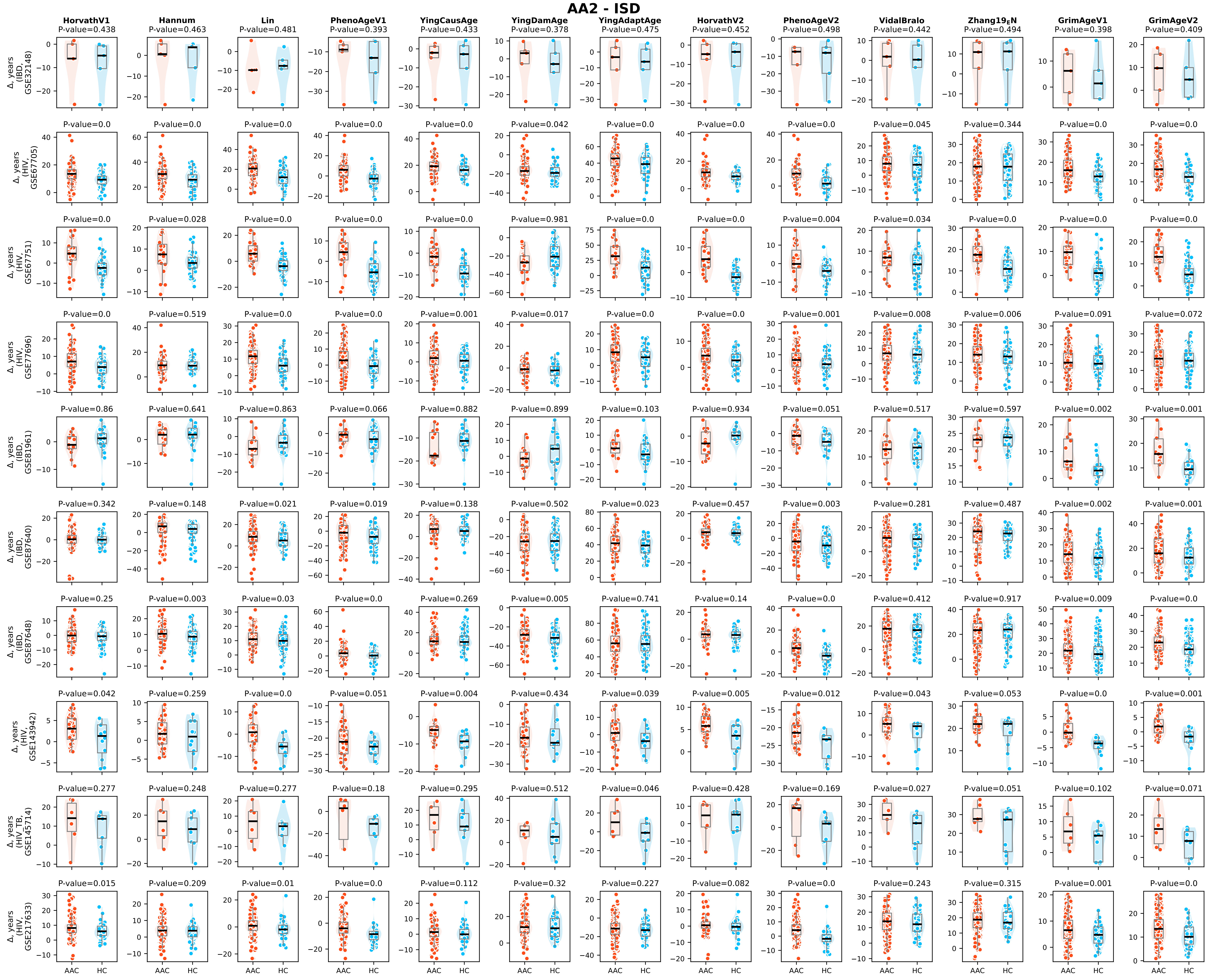

### Figure_A6_AA2-LUD.pdf

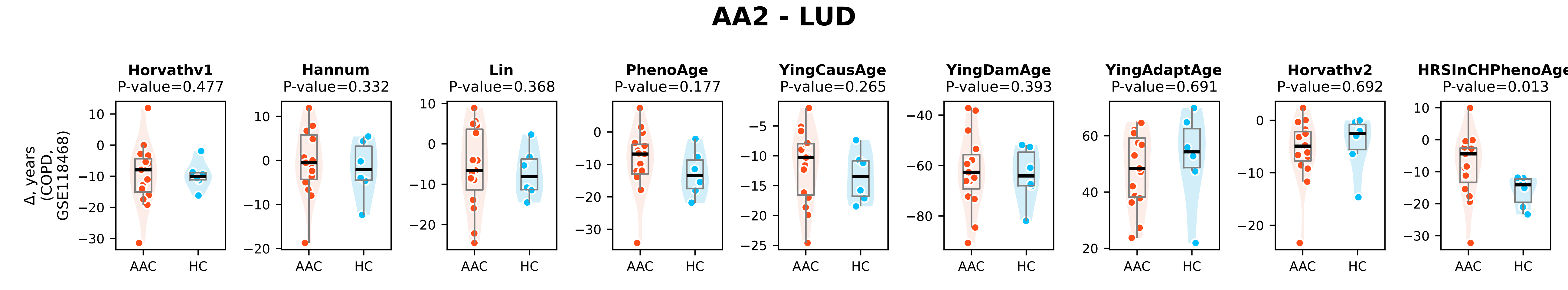

### Figure_A6_AA2-MBD.pdf

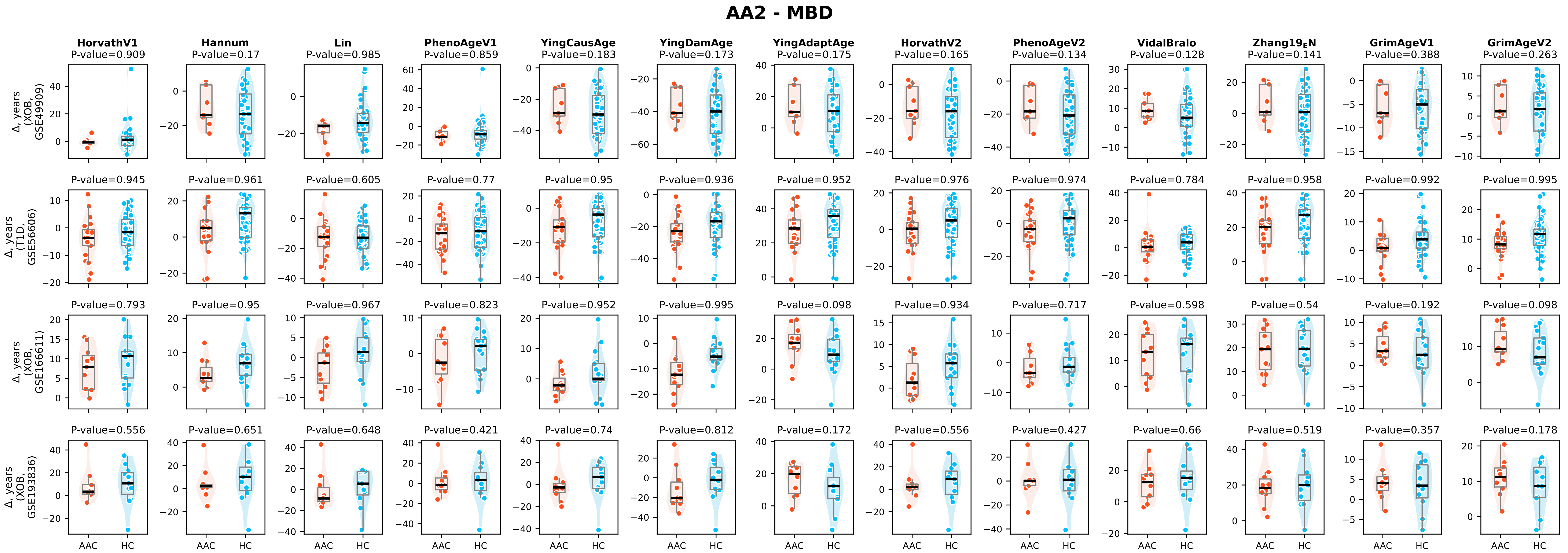

### Figure_A6_AA2-MSD.pdf

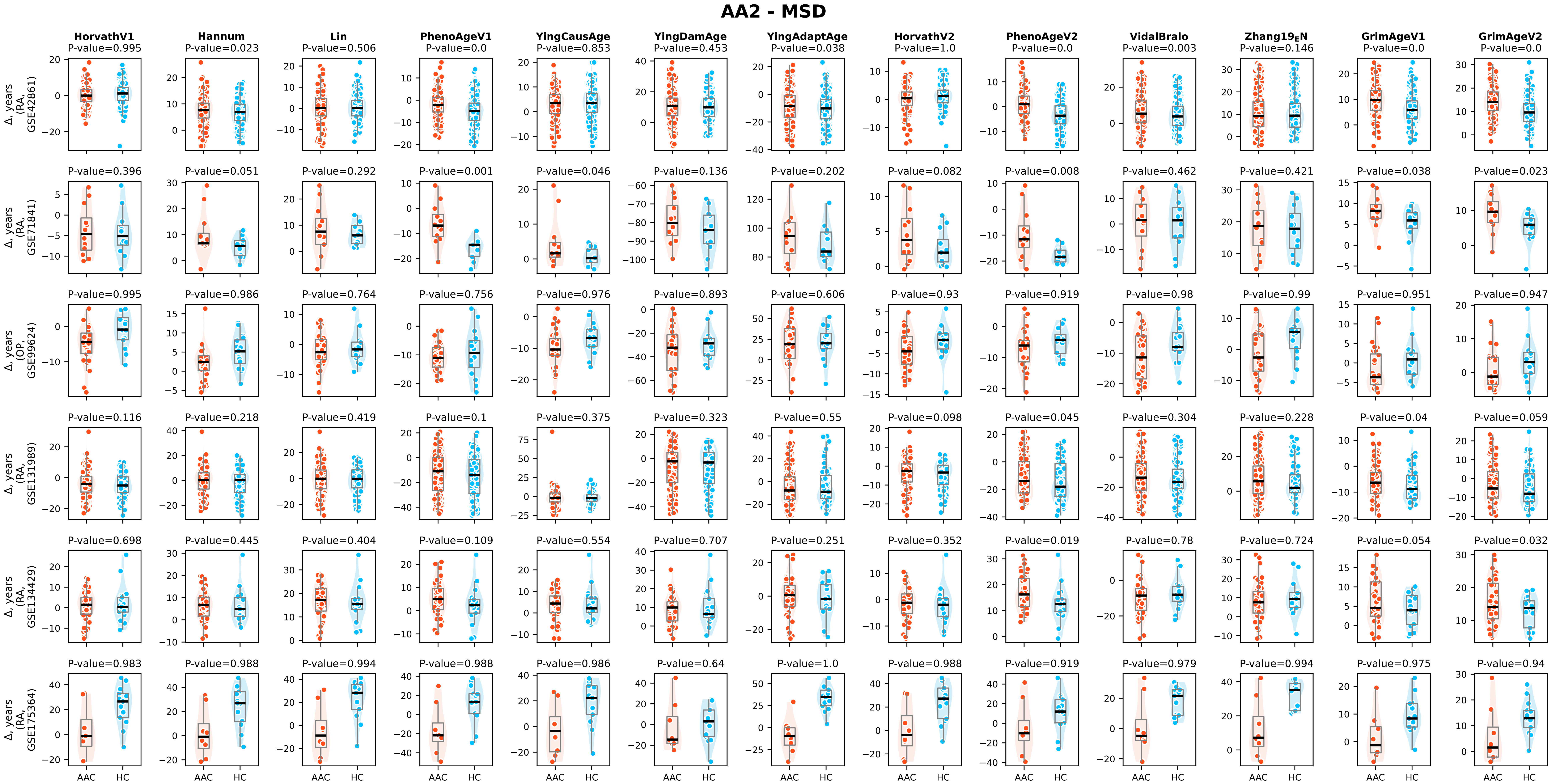

### Figure_A6_AA2-NDD.pdf

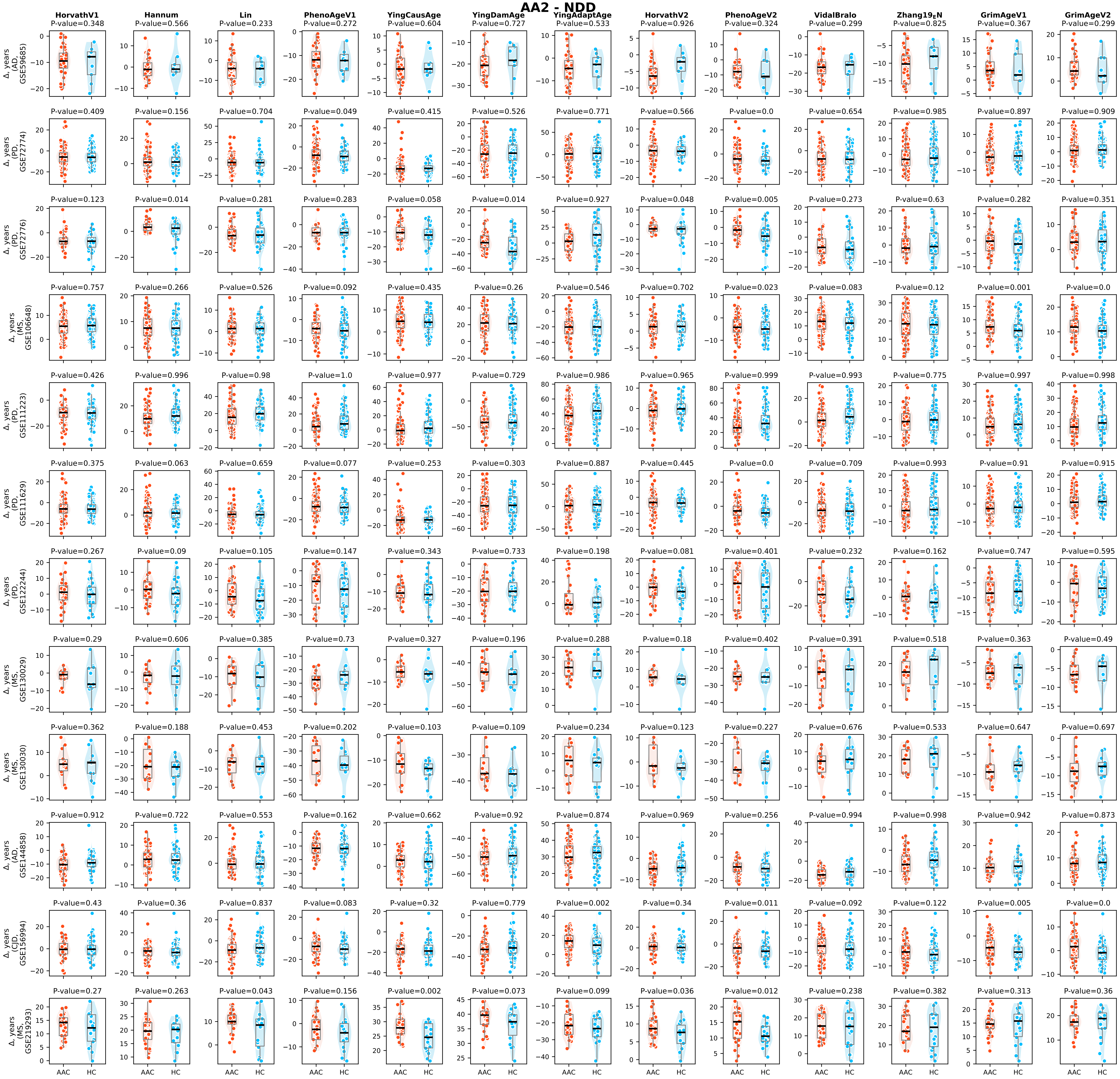

### Figure_A6_AA2-PGS.pdf

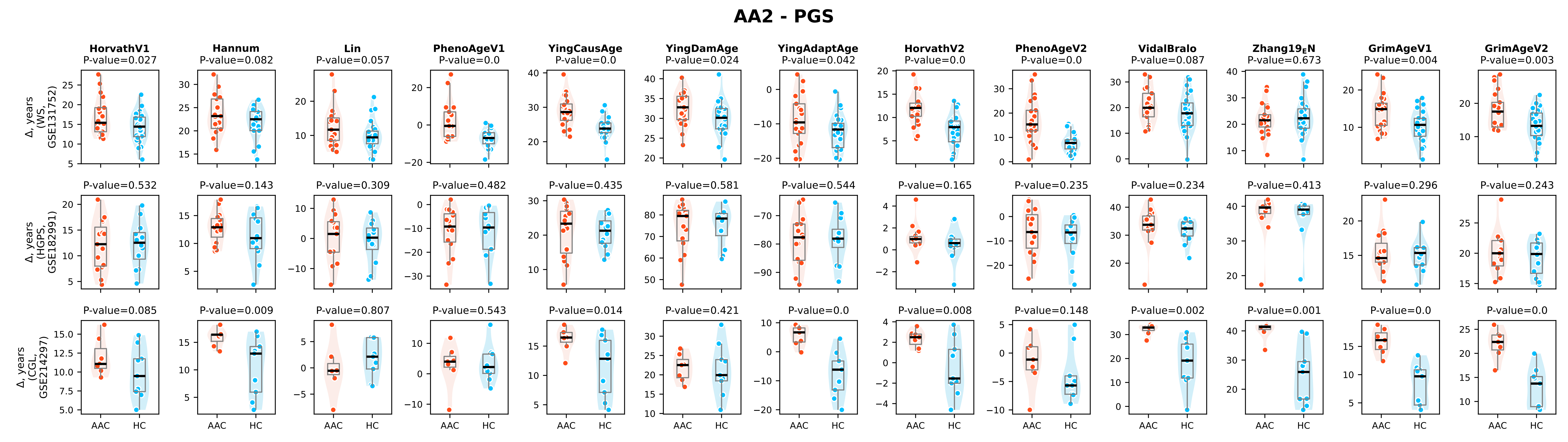

### Figure_A6_AA2-RSD.pdf

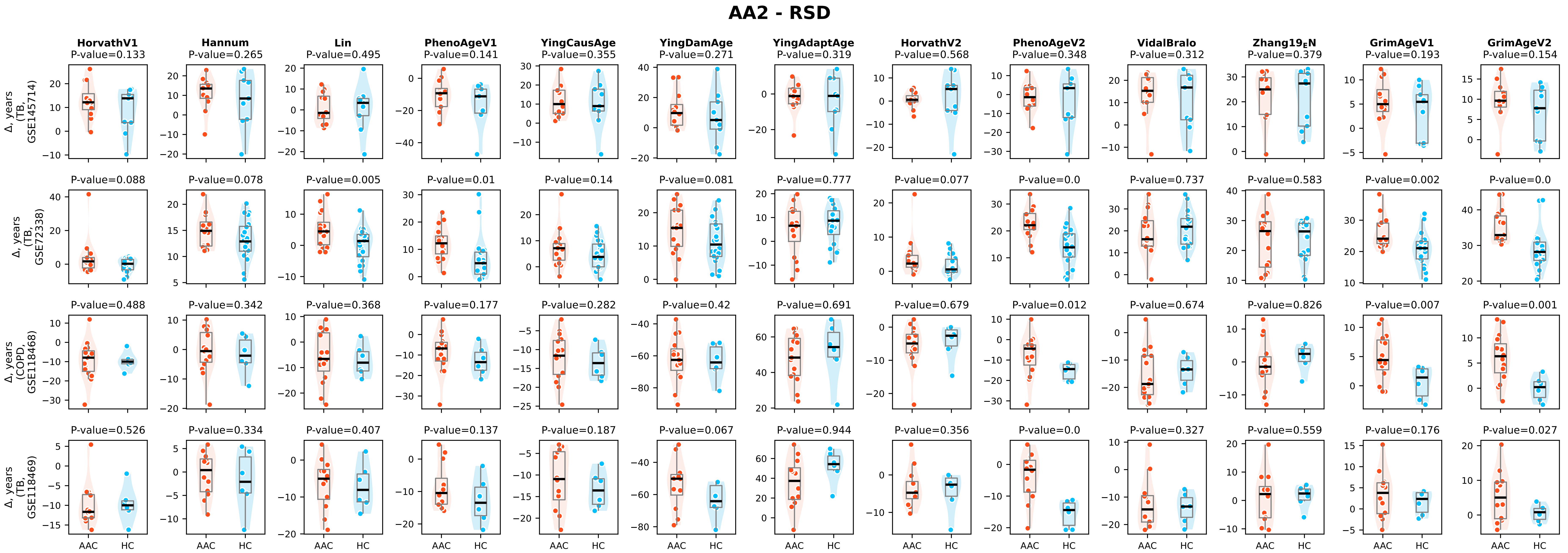

### Figure_A7_AA1-CVD.pdf

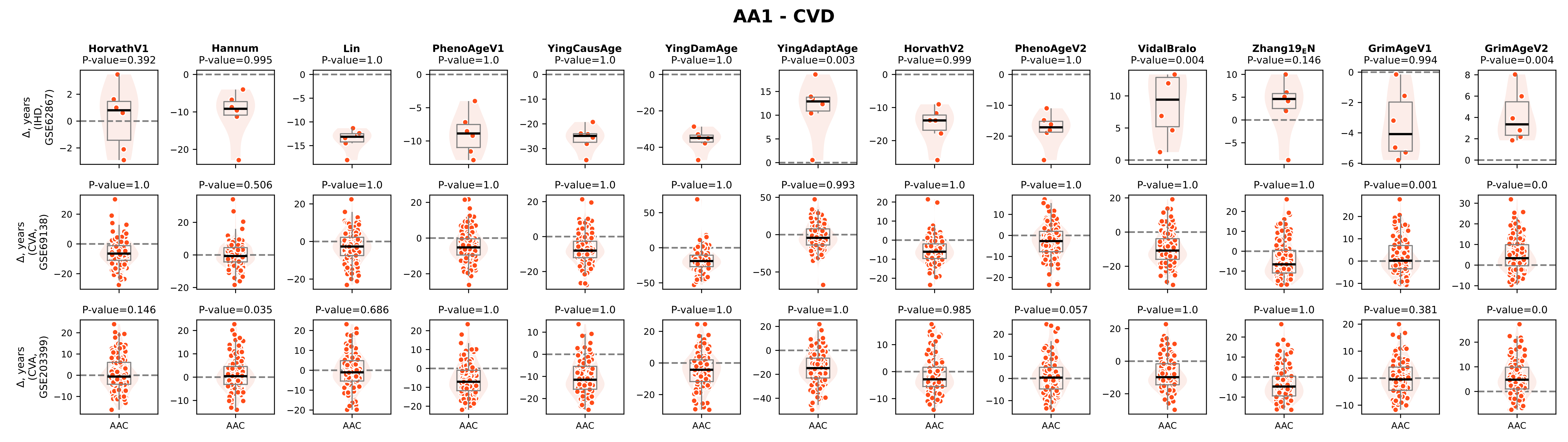

### Figure_A7_AA1-ISD.pdf

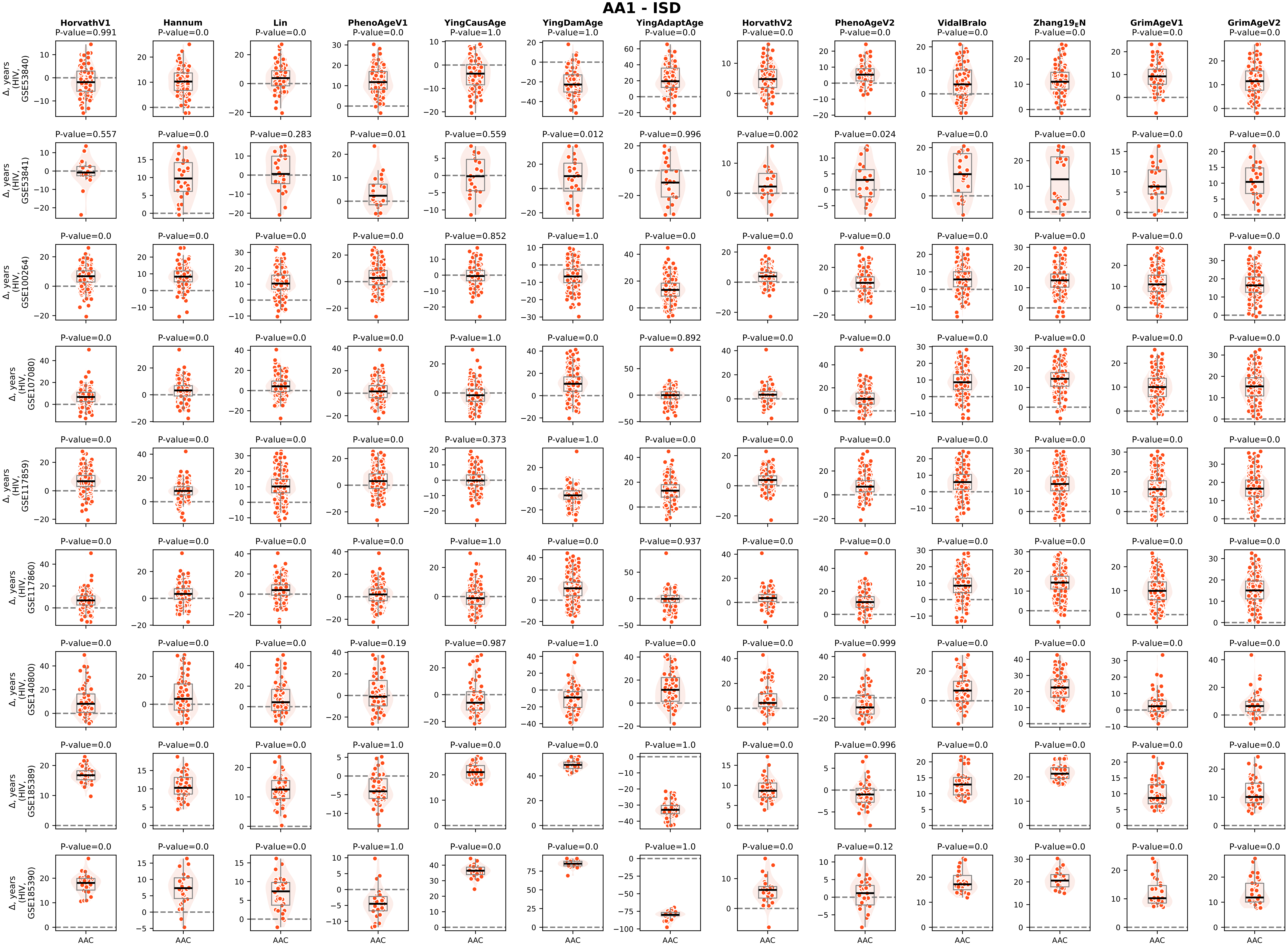

### Figure_A7_AA1-MBD.pdf

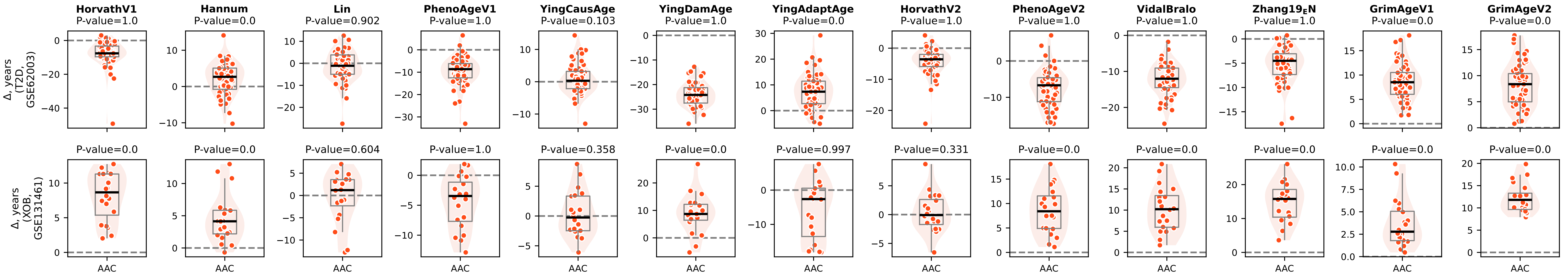

### Figure_A7_AA1-MSD.pdf

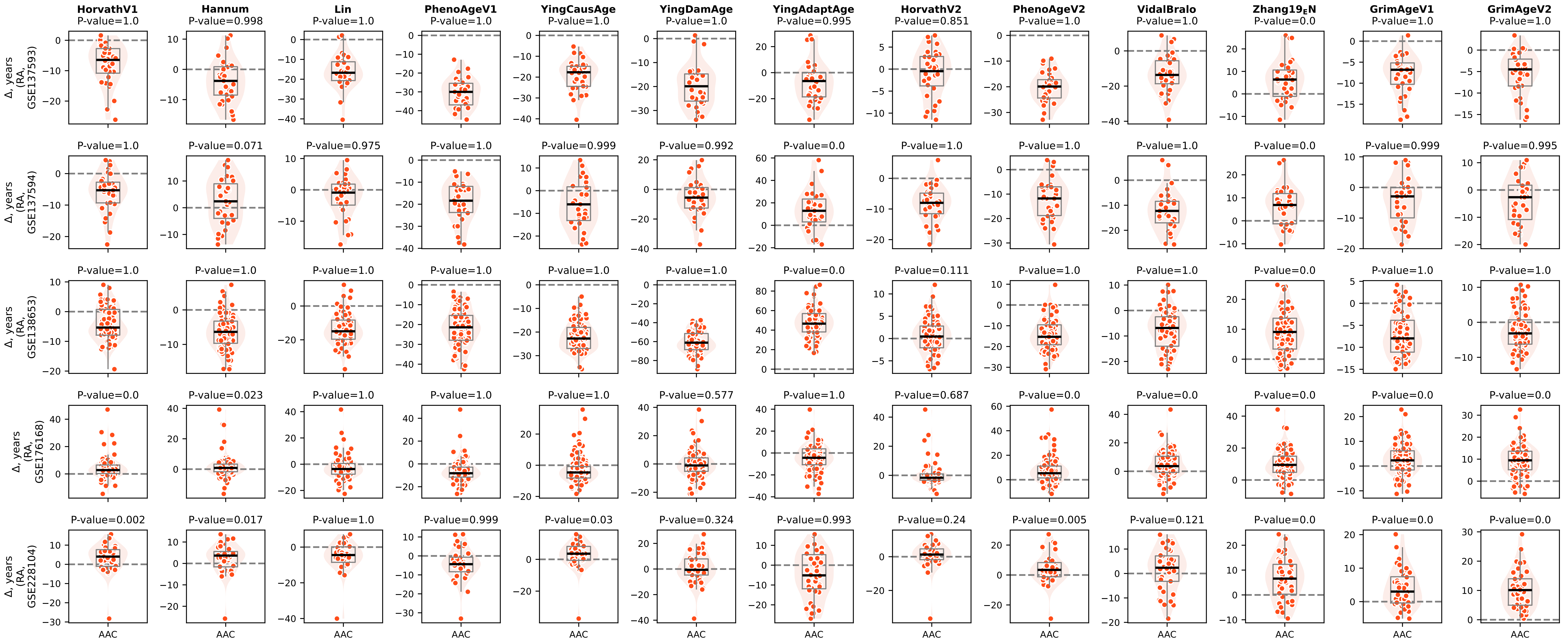

### Figure_A7_AA1-NDD.pdf

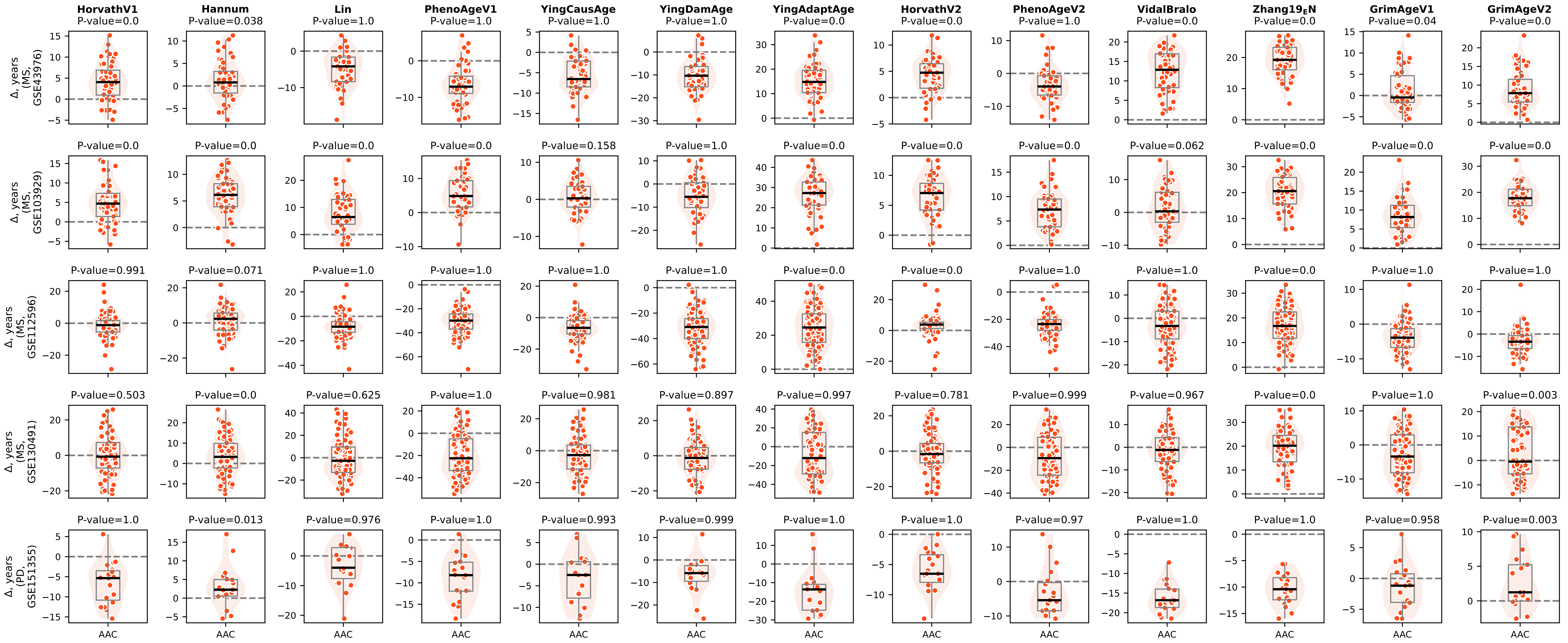
